# Differential effects of chronic restraint stress on two active avoidance tasks in rats

**DOI:** 10.1101/2025.02.28.640742

**Authors:** A López-Moraga, M. De Ceuninck, Y. Van der Heyden, L. Vercammen, R. Palme, B. Vervliet, T. Beckers, L Luyten

**Affiliations:** Centre for the Psychology of Learning and Experimental Psychopathology, Faculty of Psychology and Educational Sciences, KU Leuven, Leuven, Belgium; Leuven Brain Institute, KU Leuven, Leuven, Belgium; Laboratory of Biological Psychology, Faculty of Psychology and Educational Sciences, KU Leuven, Leuven, Belgium; Experimental Endocrinology, Department of Biological Sciences and Pathobiology, University of Veterinary Medicine Vienna, Vienna, Austria

**Keywords:** stress, active avoidance, corticosterone, rat, extinction

## Abstract

Although avoidance can serve an adaptive function in daily life, excessive or persistent avoidance can form a debilitating symptom of anxiety-related disorders. The transition from adaptive to maladaptive avoidance remains poorly understood, but stress is a potential contributing factor. We investigated the effects of chronic restraint stress on two active avoidance procedures: two-way active avoidance (2WAA) and platform-mediated avoidance (PMA). Whereas 2WAA entails a low-cost response, avoidance in the PMA task comes with a cost, i.e., no access to food. We hypothesized that chronic restraint stress would hinder avoidance acquisition in the 2WAA task, but increase avoidance acquisition in the PMA task. In two experiments, male and female rats underwent either chronic restraint stress or a control procedure. In Experiment 1, all rats (N = 31) were then trained in a 2WAA acquisition and extinction procedure, in two contexts. Stressed rats showed significantly reduced avoidance acquisition, while extinction was unaffected. In Experiment 2 (N = 32), stressed rats and controls were trained in a PMA acquisition and extinction procedure. Contrary to our hypothesis, we did not find effects on avoidance acquisition, although we found group and sex differences in lever press suppression. All rats gradually extinguished defensive behaviors during extinction. Overall, chronic restraint stress had limited effects on PMA, but significantly impaired avoidance acquisition in the 2WAA task without affecting its extinction. These divergent effects may relate to differences in response cost or differences in safety of the context (i.e., a permanent safe area in PMA, but not in 2WAA).

**Highlights:**

- Chronic restraint stress impaired acquisition of two-way active avoidance
- Chronic restraint stress did not impact acquisition of platform-mediated avoidance
- Chronic restraint stress did not impact extinction of avoidance in either task
- Minor sex differences were observed across both tasks

## Introduction

Although avoidance can serve an adaptive function in daily life, excessive or persistent avoidance can form a debilitating symptom of anxiety-related disorders. For example, staying away from dark alleys at night in large cities may be an adaptive avoidance response, because it thwarts potential threats and ensures survival. However, always staying in and avoiding even generally safe neighborhoods is arguably maladaptive, because an objectively harmless situation is treated as if it were dangerous (Arnaudova et al., 2017).

The specific factors underlying the transition from adaptive to maladaptive avoidance are not fully understood, but chronic stress is a key candidate (Daviu et al., 2019; López-Moraga et al., 2022; McEwen, 2004). Stressors activate the hypothalamic-pituitary-adrenal axis and trigger the release of glucocorticoids and noradrenaline. These hormones are involved in short-term adaptive responses to threats and in long-term responses, including effects on learning and memory. For instance, both glucocorticoids and noradrenaline enhance the consolidation of emotionally arousing experiences and have been involved in regulating memory specificity, flexibility and accuracy (Schwabe et al., 2022). Healthy responding to acute stressors does not only imply rapid glucocorticoid synthesis, but also a quick termination of the stress response to limit the effects of these hormones, as prolonged responses that persist beyond the initial stressor can lead to disease (McEwen and Akil, 2020).

The stress response is highly conserved between rodents and humans, allowing to study chronic stress and its effects in animal models (Watson et al., 2016). Gaining a deeper understanding of the potential dysregulation of avoidance by stress may help us better comprehend the pathophysiology of anxiety-related disorders and develop new interventions (Bale et al., 2019). Clinical and experimental observations suggest that prior stress may influence avoidance, but their relationship is not unequivocal (Patterson et al., 2019; Roelofs et al., 2005; Weaver et al., 2020). Clinical findings indicate that uncontrollable or unpredictable stress leads to increased avoidance (Hancock and Bryant, 2018; Zinbarg et al., 2022). Conversely, learned helplessness, which is typically obtained by extended exposure to strong stressors, is characterized by poor avoidance or escape responses (Maier, 2001).

Our recent review of the rodent literature regarding the effects of stress on subsequent avoidance tasks revealed conflicting results (López-Moraga et al., 2022). Concerning the effects of chronic stress on active avoidance, we found that chronic restraint stress typically impairs the acquisition of active avoidance in male rats (Bravo et al., 2009; Dagnino-Subiabre et al., 2005; Ulloa et al., 2010) and female rats (Gamaro et al., 1999). However, in other studies, different stress induction procedures (chronic cold swim stress, shock and variable stress) did not affect active avoidance (Wakizono et al., 2007; Weiss et al., 1975), or increased active avoidance acquisition (with chronic cold temperature stress; Hata et al., 1989). There are multiple possible reasons for these conflicting findings, including differences in the strain, sex, and age of subjects, and in the type and duration of the stress induction procedure (López-Moraga et al., 2022).

All studies mentioned above did use the same type of avoidance task, i.e., two-way active avoidance (2WAA). This task is conducted in a shuttle box with two compartments, between which the animal can freely move. The rodent first learns the association between a warning signal (e.g., a tone) and a following aversive event (e.g., a foot shock). Subsequently, it learns that if it shuttles to the other compartment during the tone, the tone is terminated and the foot shock is omitted. This way, rodents learn to perform conditioned avoidance responses (Choi et al., 2010). Inherent to the 2WAA procedure, the animal repeatedly needs to return to a place where it has received a shock before, and therefore does not have a permanent safe place (Diehl et al., 2019).

Over the past decade, an alternative active avoidance task has been developed that provides a permanent safe space: the platform-mediated avoidance (PMA) task (Bravo-Rivera et al., 2014). In this task, rats learn that they can avoid a tone-signaled foot shock by stepping onto a platform during the tone, which is a location where the animal is always safe from shocks. However, stepping onto the platform does come at the expense of lever pressing for food, which constitutes an approach-avoidance conflict because the rats are food-deprived. This feature of the PMA task, which is not present in the 2WAA task, may model an important component of decision processes in individuals with clinical anxiety, who often exhibit avoidance behaviors that result in the loss of positive outcomes or potential rewards (Diehl et al., 2019). In humans, emerging evidence suggests that acute and chronic stress increase costly avoidance in anxious individuals (Vogel and Schwabe, 2019; Weaver et al., 2020). To the best of our knowledge, the effects of chronic stress on costly avoidance in rodents (e.g., in the PMA task) have not yet been investigated.

The 2WAA and PMA tasks differ in several key aspects, as mentioned above. Therefore, it is possible that chronic stress has differential effects on platform-mediated avoidance, which involves costly avoidance, compared to two-way active avoidance, which entails a low-cost avoidance response. In a first experiment (n = 31), we hypothesized that chronic restraint stress would impair active avoidance acquisition in the 2WAA task, in line with prior studies using chronic restraint stress induction. We expected that it would also impair extinction of avoidance, which has remained uninvestigated thus far, but bears relevance to disease progress and treatment. In contrast, for the second experiment (n = 32), using a PMA task, we hypothesized based on preliminary observations in anxious individuals (Vogel and Schwabe, 2019; Weaver et al., 2020) that stressed rats would show more avoidance than control rats during acquisition and the first sessions of extinction.

## Methods

### Preregistration and data availability

Designs, procedures, sample sizes, and analysis plans were registered on the Open Science Framework (OSF) prior to the start of data collection: https://osf.io/qm6y7/. Data and scripts can also be found on OSF.

### Subjects

All experiments were performed in accordance with Belgian and European laws (Belgian Royal Decree of 29/05/2013 and European Directive 2010/63/EU) and the ARRIVE 2.0 guidelines (Percie du Sert et al., 2020) and were approved by the KU Leuven animal ethics committee (project license number: 177/2023). Based on previous reports that investigated the effects of chronic restraint stress on active avoidance acquisition in male rats (Dagnino-Subiabre et al., 2005; Ulloa et al., 2010), we calculated an effect size (Cohen’s d) of 2.66. Given this effect size was large and could be a result of publication bias, we decided to set power to 95% and use a Cohen’s d of 1.3, half of the published d, to calculate our sample size. Using pwr in R we found that n = 16 per group would be enough to achieve our desired power, resulting in a sample size of 32 rats per experiment.

Both experiments were conducted using 7-8-week-old female (180-200 g at arrival) and male (250-280 g at arrival) Wistar rats (Janvier Labs, Le Genest_-_Saint_-_Isle, France). Animals arrived in our facility 4 (Experiment 1) or 5 days (Experiment 2) before the start of the experiment. Animals were housed in groups of 4. The housing room has a 12-hour light-dark cycle (lights on at 7 am), and experiments were performed between 8 am and 6LJpm.

Animals of the stress group were housed in the same room, but in a different ventilated cabinet than animals of the control group. The cages had bedding and cage enrichment in the form of a red polycarbonate tunnel hanging from the top of the grid. Water was available ad libitum for the entire experiment, except during behavioral testing. For Experiment 1, food was provided ad libitum, except during behavioral testing (see Figure 1a). For Experiment 2, food was provided ad libitum except one day before and during lever press training, immediately after the sucrose preference test, and during all avoidance and extinction days in the platform-mediated avoidance task (see Figure 1b). Animals in this experiment were handled for 2 days before the start of lever press training.

**Figure 1.**
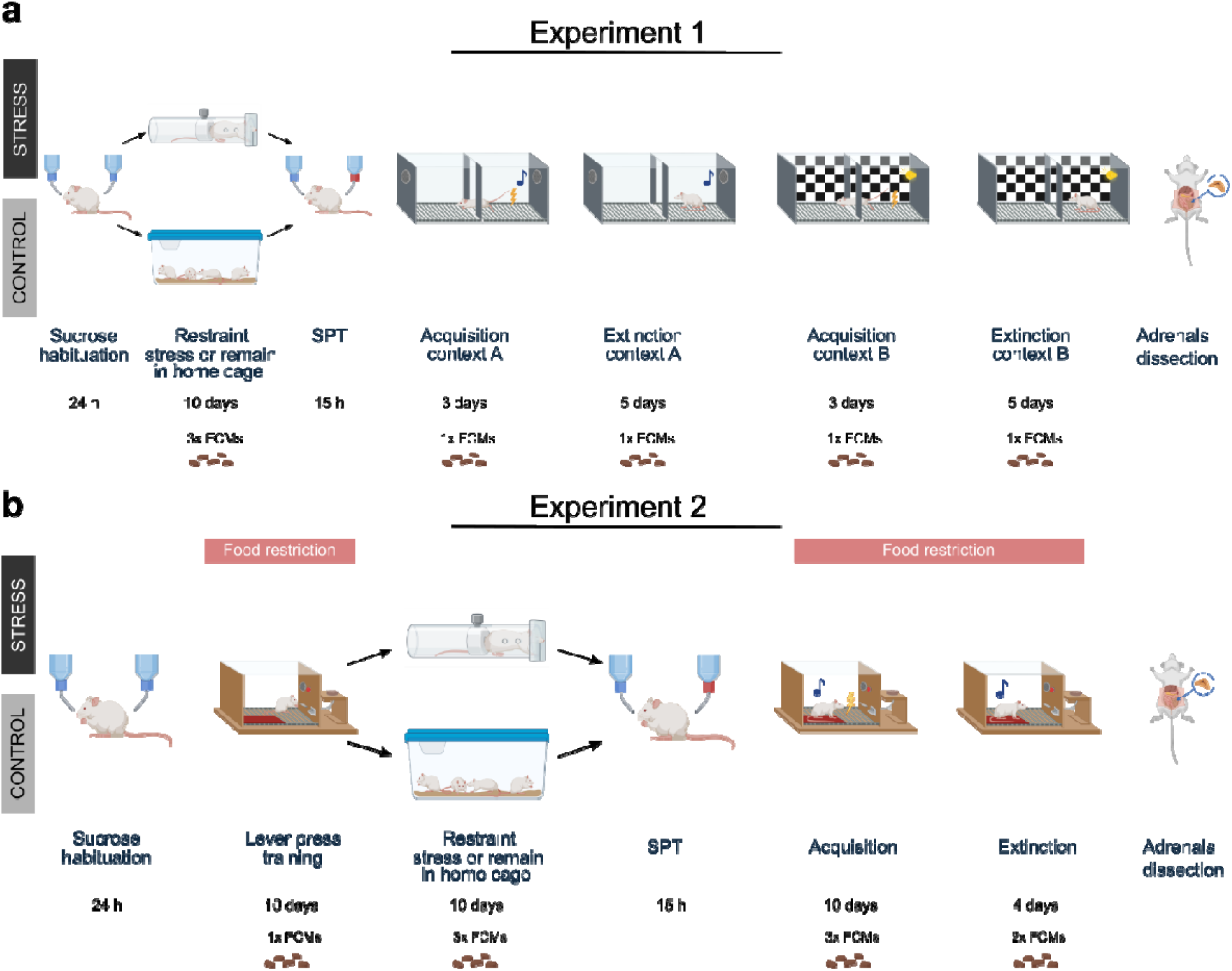
The experimental timeline of Experiment 1 (**a**) and Experiment 2 (**b**). SPT: sucrose preference test. FCMs: fecal corticosterone metabolites.

In Experiment 1, one male rat from the stress group died unexpectedly on day 6 of the stress induction procedure and was excluded from all analyses, resulting in a total sample of 31 animals in this experiment.

### Body weight measurement

Body weight was measured after every stress induction session (control animals were weighed in their housing room while the others were in the restraint stress holder) and after every behavioral session.

Previous research has shown that restraint stress negatively affects body weight gain (Jeong et al., 2013). As previous authors have reported (Dagnino-Subiabre et al., 2005; Ulloa et al., 2010), monitoring changes in body weight can serve as an effective manipulation check for stress induction. Weight change over the stress induction period was calculated by taking the weight of the rat on a given day and dividing it by its baseline weight. For Experiment 1, the baseline weight was obtained 1 day before the start of the stress induction period. For Experiment 2, the baseline weight was defined as the weight on the first day of the stress induction, due to food restriction until the day before the start of stress induction.

### Sucrose preference test

The sucrose preference test has been used to evaluate the effects of chronic stress (Katz, 1982; Willner et al., 1987). Two days before the start of stress induction, rats from Experiment 1 were habituated to consume a 1% sucrose (Acros Organics) solution for 24 h, with two bottles of a 1% sucrose solution placed in their home cage and no access to regular water. Rats from Experiment 2 underwent this sucrose habituation one day before the lever press training started. For both experiments, one day after the stress induction period, all rats were housed separately for 15 hours in smaller, individual cages and subjected to a sucrose preference test. Food was provided ad libitum, and water was provided in two bottles: one with tap water and one with the 1% sucrose solution, from 18:30 to 9:30 (Berrio et al., 2024; Liu et al., 2018). Given that rats have a choice between 1% sucrose in tap water and pure tap water, a reduced preference for the sweet solution compared to tap water may be an indication of anhedonia-like behavior and has been observed after chronic stress induction in rodents (Berrio et al., 2024; Mao et al., 2022). The position (left or right) of the two different bottles was counterbalanced across cages. After the test, rats were again housed in their usual group home cages. The weight change of the bottles was used as a measure of intake. The sucrose preference ratio was calculated using the following formula: Sucrose Preference ratio = Sucrose solution intake (g) / [Water intake (g) + Sucrose solution intake (g)].

### Chronic restraint stress induction

Rats from the stress group were brought to a room close to the behavioral testing rooms on a cart in groups of two cages (one with 4 males and one with 4 females), before the start of each stress induction session. Rats were placed in a cylindrical restrainer (males: #51335; females: #51336, Stoelting Europe, Dublin, Ireland) for 2 h daily for 10 consecutive days.

Half of the rats underwent restraint before noon and the other half in the afternoon. During the restraint sessions, the cylinders were placed in individual cages (type III) with a metal grid on top. For purely exploratory purposes, an ultrasound microphone (Avisoft-Bioacoustics CM16/CMPA, Glienicke, Germany) was attached with zip ties to the metal grid to record the occurrence of any aversive 22-kHz ultrasonic vocalizations (USV) emitted by the male rats (Wöhr & Schwarting, 2013). After each stress induction session, the cages, restrainers, and surfaces of the room were cleaned with Ecover All Purpose Cleaner, Lemongrass & Ginger (5%). Control rats remained undisturbed in their home cages, handled only to perform daily weighing and to obtain fecal samples.

### Two-way active avoidance (Experiment 1)

The effects of prior chronic restraint stress on 2WAA acquisition and extinction in two contexts were tested in 15 male and 16 female rats (see Figure 1a).

### Apparatus

Four identical operant chambers (28.7 width, 24.13 cm depth and 28.7 cm height; ENV-010MXL, Med Associates Inc, Vermont, USA) were used simultaneously and were enclosed in sound-attenuating cubicles (ENV-020M, Med Associates). Each operant chamber was divided into two identical compartments by custom doorway panels. Behavior was recorded with an infrared IP camera (Foscam C1, Shenzhen, China) attached to the ceiling of the sound-attenuating cubicle. Tones were generated by a multiple tone generator (ENV-223, Med Associates) and speaker (ENV-224AM-3, Med Associates). The shock floor (ENV-005FPU-R, Med Associates) of each compartment consisted of 19 stainless steel rods 0.47 cm in diameter and spaced 1.59 cm apart, center to center. Each side of the compartment contained four independent infrared photobeams (ENV-256-8S, Med Associates) to track where the rat was located. All experimental sessions were programmed using Trans™ editor and were carried out by MED-PC™ software.

### Acquisition and extinction

Rats were tested in batches of four. The order of testing was counterbalanced across experimental group and sex. All animals first underwent 3 days of avoidance acquisition and 5 days of avoidance extinction in context A, and then 3 days of acquisition and 5 days of extinction in context B.

#### Context A

The shuttle box was not illuminated and was cleaned with a pine-scented cleaning product (econom Allesreiniger dennenfris, 5%). The CS was a tone with an intensity of 75 dB and a frequency of 3kHz. First, rats underwent 3 days of avoidance training in context A (Vercammen et al., 2025). On each day, avoidance training started with a 5-min acclimation period, whereafter 30 CSs of maximum 20 s were presented, with an ITI averaging 70 s (40 s to 100 s). Each CS co-terminated with a foot shock US of maximum 10 s and an intensity of 0.4 mA. All rats had the possibility to avoid or escape the US by shuttling to the opposite compartment during or right after the tone, respectively. If the rat shuttled to the other compartment during the presentation of the CS, the US was avoided, the CS was terminated and the trial was labeled as an avoidance response. If the rat shuttled to the other compartment during US presentation, the US was terminated and the trial was labeled as an escape response. After 3 days of avoidance training, rats went through extinction training for 5 days, where each day started with a 5-min acclimation period, whereafter 24 CSs of the same characteristics as above were presented per day, without foot shock. Note that for video synchronizing purposes, the onset of the tone CS used in context A was accompanied by a dim 2-s light cue (a partially covered house light at the center of the back wall) oriented towards the camera.

#### Context B

Forty-eight hours after the last extinction session in context A, all rats were again subjected to 3 days of avoidance training and 5 days of extinction, however this time in a different context B, to further evaluate the longevity of any effects of the chronic restraint stress induction. To change the context, a checkerboard pattern was added to the outside of the long transparent sides of the box. The inside of the box was dimly lit using two houselights (3.7 lux, measured at the center of each compartment’s grid). The scented cleaning product was changed to Instanet (5%). And lastly, the CS used in context B was a light stimulus (round yellow LED, 3 cm above the grid floor, MedAssociates, ENV-221M-LED, 40 lux) with a maximum duration of 20 s.

### Platform-mediated avoidance (Experiment 2)

The effects of prior chronic restraint stress on PMA acquisition and extinction were tested in 16 male and 16 female rats (see Figure 1b).

### Apparatus

Eight identical operant chambers (30.5 cm width, 25.4 cm depth and 30.5 cm height; Rat Test Cage, Coulbourn Instruments, Pennsylvania, USA) were used simultaneously and were enclosed in individual sound-attenuating boxes. A 12.8 cm by 15.2 cm platform made of a non-transparent red plastic 3-mm sheet was used in all sessions. It approximately covered ¼ of the grid floor and was placed in the corner opposite to the lever. Experiments were conducted using Graphic State 4 (Coulbourn Instruments) with the house light off. Behavior was continuously recorded during the experimental sessions using an IP camera (Foscam C2M, Shenzhen, China).

### Lever press training

Lever press training consisted of 45 min of training for 10 consecutive days. Rats were gradually shaped to lever press for grain-based pellets (Grain-Based Dustless Precision Pellets® for Rodent, Bio-Serv, Frenchtown, NJ, USA). The first training phase consisted of one magazine training session with the lever retracted, during which food pellets were delivered at fixed 2-min intervals. The second phase of lever press training consisted of magazine training with the lever extended, during which food pellets were delivered at fixed 2-min intervals, supplemented with direct food pellet delivery upon each lever press. This phase lasted a minimum of 1 day and required at least 1 lever press/min on average to pass on to the next phase. If there were no lever presses during the first half of the third day of the training phase, hand shaping was performed. The next phases consisted of lever press training on increasing variable ratio (VR) schedules, progressing from VR 3 through VR 5 and VR 15 to VR 30, where pellets were delivered after a variable number of lever presses, with an increasing average reinforcement criterion per session. For VR 3, a criterion of at least 3 lever press/min was set to pass to the next schedule, a criterion of at least 5 lever press/min for VR 5, a criterion of at least 10 lever press/min for VR 15, and a criterion of at least 15 lever press/min for VR 30. The final training phase consisted of training on a variable interval (VI) schedule of reinforcement averaging 30 seconds (VI 30). The first lever press after a variable refractory period averaging 30 s yielded pellet delivery.

Rats were included if they met at least one of two preregistered criteria: 10 lever press/min after averaging the first 10 min of all lever-pressing training sessions in VI 30 or 10 lever press/min after averaging the complete duration of all lever-pressing training sessions in VI 30. All rats met at least one of the two learning criteria; therefore all 32 rats were included.

### Acquisition and extinction

Acquisition of platform-mediated avoidance started 1 day after the end of the sucrose preference test. Avoidance acquisition lasted for 10 days in accordance with previous research (Bravo-Rivera et al., 2015, 2014; Diehl et al., 2018; Martínez-Rivera et al., 2019). Each day, 9 30-s CSs were presented, with an intertrial interval (ITI) averaging 180 s (150 s to 210 s). This ITI was also used in subsequent sessions. The CS consisted of a pure 3-kHz tone co-terminating with a 2-s, 0.4-mA foot shock US, which was delivered through the grid floor. The plastic platform in one corner constituted a safe space where the rat would not experience the foot shock during US presentation. The experiment ended with four daily extinction training sessions in accordance with previous research (López-Moraga et al., 2024). Each extinction session consisted of the 30-s CS being presented nine times, as in avoidance acquisition, but without US delivery.

### Fecal corticosterone metabolites

At several time points, we collected fecal boli to determine fecal corticosterone metabolites (FCMs) as a proxy for systemic corticosterone levels. This non-invasive method has previously been validated in rats (Lepschy et al., 2007; Palme, 2019). In Experiment 1, fecal boli were collected at the following time points (Figure 1a): immediately before sucrose habituation, on days 2, 6 and 10 of restraint stress, day 2 of 2WAA acquisition in context A, day 2 of 2WAA extinction training in context A, day 2 of 2WAA acquisition in context B and day 2 of 2WAA extinction training in context B. In Experiment 2 (Figure 1b), fecal boli were collected on day 10 of lever press training, days 2, 6 and 10 of restraint stress, days 2, 6 and 10 of platform-mediated avoidance acquisition and days 2 and 4 of extinction training. Fecal boli were collected from the restraint cylinder or behavioral testing setup immediately after the end of the session. On days 2, 6 and 10 of the stress induction period, fecal boli of the control group were collected during their daily weighing measurements.

Fecal boli were immediately stored atLJ-70°C until processing. Feces were dried in an oven at 70 °C (COOK-IT Mini Oven 21L, Media Evolution B.V, Eindhoven, The Netherlands) for 4 h. Afterwards, they were reduced to powder using a mortar and pestle, and 0.05 g was weighed and mixed with 1 ml of 80% methanol (Sigma-Aldrich, Schnelldorf, Germany) in deionized water. Samples were placed in a multivortex (Thermomixer comfort, Eppendorf, Hamburg, Germany) at 21 °C for 30 minutes at 1400 rpm and then centrifuged (Centrifuge 5424, Eppendorf) for 10 minutes at 2500 g. Next, 0.5 ml of the supernatant was collected and stored at −70 °C until analysis. Samples were shipped and a group-specific 5α-pregnane-3β,11β,21-triol-20-one enzyme immunoassay, which measures a group of corticosterone metabolites with a 5a-3ß,11ß-diol configuration (for details of the assay see: Touma et al., 2003), was performed in the Laboratory of Experimental Endocrinology at the University of Veterinary Medicine Vienna as previously described (Lepschy et al., 2010, 2007).

### Adrenal gland dissection

One to two days after the last behavioral test session, rats were injected intraperitoneally with an overdose of sodium pentobarbital (200 mg/ml solution for injection, Vetoquinol B.V.). In Experiment 1, the adrenal glands were extracted immediately after pentobarbital injection. In Experiment 2, rats were first transcardially perfused with 1% phosphate-buffered saline followed by 4% paraformaldehyde solution before dissecting the adrenals. Any visible adipose tissue was surgically removed, and the glands were weighed. Chronic stress can cause the enlargement of the adrenal glands, resulting in an increased organ weight (Ulrich-Lai et al., 2006), making adrenal gland-to-body weight ratio a chronic stress manipulation check that other authors previously used (Dagnino-Subiabre et al., 2005). Adrenal gland-to-body weight ratio was calculated by averaging the weight of the two adrenal glands and dividing this by the body weight of the rat prior to euthanasia.

### Behavioral outcomes

#### Experiment 1

Avoidance responses, avoidance latencies, escape latencies and crossings were measured via the infrared beams in each compartment and collected by MED-PC software. Additionally, freezing during the 5-min acclimation period was scored from video recordings. Freezing was defined as full immobility except for the minimal movements associated with breathing.

Scoring was done manually by the experimenter, who was blinded to group allocation during scoring. The experimenter could not be blinded to the phase of the experiment (avoidance training or extinction, and context A or B) and sex during scoring, because this information is visible on the videos.

#### Experiment 2

Lever presses were recorded using Graphic State 4 software (Coulbourn Instruments, Pennsylvania, USA) and further processed using an in-house R script available on OSF. Lever presses produced 1 min before the tone (pretone) and during the tone were aggregated per block (each block consisting of 3 tones), after which the following formula was applied to calculate the suppression of lever pressing: (pretone rate - tone rate)/(pretone rate + tone rate) x 100 (Quirk et al., 2000). The obtained values ranged from no suppression of lever pressing (0%) to full suppression of lever pressing (100%). For blocks in which pretone and tone values were both 0, the rat’s suppression was substituted by the mean group average for that block.

Time spent on the platform (i.e., avoidance) and freezing during CS presentations were scored using a custom automated scoring pipeline that relied on DeepLabCut (DLC) and SimBA (see below). To check its validity, we compared 16 videos (8 from acquisition and 8 from extinction sessions) analyzed using our automated scoring pipeline with manual scoring by trained observers and obtained an intraclass correlation coefficient (ICC) of 0.91, 95% CI [0.88, 0.94] for the avoidance measurement and an ICC of 0.88, 95% CI [0.83,0.91] for the freezing measurement (see Validation folder on OSF). Given the high ICC, we proceeded with automated scoring only.

We processed our videos using DLC and SimBA to obtain speed traces of each rat in cm/s. As previously described (Gruene et al., 2015), we considered a discrete darting bout when an animal moves with a minimum velocity of 23.5 cm/s during any CS presentation. An animal is considered a darter if it exhibits at least one dart between the 3rd CS-US pairing and the 7th CS-US pairing of the first avoidance training session (Gruene et al., 2015; López-Moraga et al., 2025).

#### DeepLabCut

We used DeepLabCut (version 2.3.9) for body part tracking (Mathis et al., 2018; Nath et al., 2019). We labeled 196 frames taken from 10 videos, each video originating from a different animal; 95% of these frames were used for training. Five body points were selected: nose, left ear, right ear, centroid, and tail base. We used a RestNet-50 neural network (He et al., 2015; Insafutdinov et al., 2016) with default parameters for 300000 training iterations. We validated with 1 shuffle and found that the test error was 4.78 pixels, and the training error was 1.88 pixels (image size: 455 x 256 pixels, downsampled from original 1280 x 720 pixels).

#### SimBA

To further analyze pose estimation data obtained using DeepLabCut, we used SimBA (Goodwin et al., 2024) to automate the analysis of different behaviors in the PMA as described previously (López-Moraga et al., 2025). We created a single animal project configuration with user-defined body points. We defined pixels per mm by measuring the width of the bottom of the fear conditioning box.

To analyze avoidance behavior, we defined the platform and adjacent wall sections as the region of interest (ROI). If the centroid body part was inside the ROI, this was considered as avoidance behavior.

To analyze darting behavior, we used the Analyze Distances/Velocities module with a 1-second time bin. We then used our own scripts to identify time bins with CS presentations (see OSF). The last 2 s of the CS presentations were excluded, as the CS co-terminated with the foot shock, and darting could be confounded with a US escape response.

We constructed a random forest classifier in SimBA to assess freezing. Eight videos were manually annotated; 22505 frames (750.17 s) were annotated with freezing present and 137638 frames (4587.93 s) with freezing absent. The start frame was defined as the first frame in which the animal stopped moving, and the end frame was defined as the last frame before the animal started moving again. We built the model with the following settings: n_estimators = 2000, RF_criterion = gini, RF_max_features = sqrt, RF_min_sample_leaf = 1, under_sample_ratio = 4, class_weights = custom: Freezing present = 2; Freezing Absent = 1, with an 80% training split. The evaluation performance of the freezing classifier had a precision score of 0.87, recall 0.70, and F1 score of 0.78 for freezing present, and a precision score of 0.92, recall 0.97, and F1 score of 0.94 for freezing absent. We set the discrimination threshold at Pr = 0.65, and a minimum duration of 500 ms. We used our own scripts to identify time bins with CS presentations to further analyze freezing instances (see OSF).

### Data analyses

Statistical analyses were preregistered, and any deviations are specified below. Data were analyzed using two-way, repeated, or mixed ANOVAs, as appropriate. Negative effect sizes of ANOVA tests (ω_p_^2^) are reported as 0 (Kroes and Finley, 2023). Post-hoc tests with multiple test corrections followed if significant main effects or interactions were found. If assumptions were violated, nonparametric statistical analyses were performed. For the nonparametric mixed RM ANOVAs, we applied an aligned rank transform ANOVA using the ARTool (Elkin et al., 2021) package from R (Version 4.1.3; R Core Team, 2022) and R Studio (Version 4.2.764; Posit Team, 2024). All other statistical analyses were performed using performance (Lüdecke et al., 2021), afex (Singmann et al., 2023), emmeans (Lenth et al., 2024) and effectsize (Ben-Shachar et al., 2024) R packages. Data were processed and plotted using the tidyverse (Wickham and RStudio, 2022) and zoo (Zeileis and Grothendieck, 2005) packages. The USV recordings were analyzed using Avisoft SASLabPro Software (Avisoft-Bioacustics, Glienicke/Nordbahn, Germany). As the initial analysis did not indicate any aversive 22 kHz calls, no further analysis of vocalizations was conducted.

### Stress induction

To evaluate the effectiveness of the chronic restraint stress induction, we assessed its effect on four variables: weight change, sucrose preference ratio, adrenal glands weight and FCM levels.

We performed a mixed ANOVA with day as a within factor, group as a between factor and weight change as a dependent variable. To analyze the sucrose preference ratio, we used a two-way ANOVA with sucrose preference ratio as the dependent variable and sex and group as between factors. Adrenal gland-to-body weight ratio was used as dependent variable in a two-way ANOVA with sex and group as factors. Differences in FCM levels (log-transformed) were analyzed with a mixed ANOVA with day (baseline, day 2, 6 and 10) as within factor, and sex and group as between factors. By omission, the inclusion of the baseline in FCMs and weight was not preregistered.

### 2WAA analyses

To evaluate avoidance behavior in Experiment 1, we conducted between-group, between-sex and within-day comparisons of number of avoidance responses during 3 days of acquisition in context A and 5 days of extinction in context A with mixed ANOVAs (Group x Sex x Day). In addition, we carried out separate analyses between groups and sex on the first day of acquisition in context A and on the first day of extinction in context A, in which we divided the 30 CSs of day 1 acquisition into 3 blocks of 10 CSs and the 24 CSs of day 1 extinction into 3 blocks of 8 CSs. Mixed ANOVAs (Group x Sex x Block) were then performed on these datasets. The same analyses were repeated for acquisition and extinction in context B. To test a secondary hypothesis, we conducted a two-way ANOVA (Sex x Group) analysis of freezing during the acclimation of day 1 acquisition in context A. Because no freezing was observed during this period, we exploratively analyzed freezing during acclimation of day 2 acquisition in context A, using a two-way ANOVA (Sex x Group) (non-preregistered analysis).

The following analyses that we conducted for Experiment 1 were not preregistered. To test for effects of chronic restraint stress on locomotion, we conducted between-group and between-sex comparisons of crossings during the acclimation period of day 1 acquisition in context A. We also conducted between-group and between-sex comparisons of number of crossings during the intertrial interval periods of acquisition in context A (Group x Sex x Day). Mean avoidance and escape latencies during acquisition in context A were tested using RM ANOVA (Group x Sex x Day). Additionally, we conducted between-group and between-sex comparisons of mean avoidance and escape latencies during acquisition in context B (Group x Sex x Day). We explored if there were differences between the first day of acquisition in context A and the first day of acquisition in context B with a mixed ANOVA (Group x Sex x Day). Similarly, we explored if there were differences between the last day of extinction in context A and the last day of extinction in context B with a mixed ANOVA (Group x Sex x Day).

### PMA analyses

To evaluate avoidance and fear-related behavior in Experiment 2, we conducted three mixed ANOVA analyses to examine the percentage of avoidance, freezing, and suppression of lever pressing behavior between stressed and control rats. We included days 1 to 3 as within-subject factors and sex and group as between-subject factors. Similarly, we performed three mixed ANOVA analyses to investigate differences in the percentage of avoidance, freezing, and suppression of lever pressing behavior across the three blocks (3 CSs per block) of extinction (days 1-2), with sex and group as between-subject factors.

In addition, we evaluated persistent avoidance behavior. A rat was considered a persistent avoider when the time on the platform during the CS was more than 50% during the first block of the third extinction session. We compared the % of persistent avoider rats in the stress group versus the control group with a Chi-square test. Additionally, we performed four two-way ANOVAs with sex and persistent avoidance as between-subjects factors to analyze differences in freezing and suppression of lever pressing during the last avoidance acquisition day and the third extinction day. Given that there were no differences between sex or group in a Chi-square test, we investigated if there were differences between persistent avoiders and non-persistent avoiders regardless of group allocation on the last day of acquisition and on the third extinction session, as preregistered.

Finally, we assessed darting behavior in this task. The classification of darters and non-darters was performed as preregistered and as reported by other authors (Gruene et al., 2015). We preregistered a comparison of the % of darter rats in females versus male rats during avoidance training using a Chi square test. We exploratorily investigated the % of darter rats in control versus stressed rats during avoidance training using a Chi square test. Given that the assumption of more than five counts per cell was violated, we used the Fisher’s exact test as well, which yielded similar results (see OSF). We preregistered that if the Chi-square test comparison between male and female rats was significant, we would conduct further analyses. Since this was not the case, we did not perform these analyses. Additionally, due to the low number of stressed darter rats, it was not possible to run further exploratory analyses comparing darters vs non-darters between the stress and control conditions.

## Results

### Experiment 1

#### Chronic restraint stress affects body weight but not sucrose preference, relative adrenal weight or fecal corticosterone metabolites

Body weight change relative to the day before stress induction or control treatment was assessed in all rats (Figure 2a). We found a significant interaction between group and day (F(3.55, 102.87) = 73.42, p < 0.001, ω_p_^2^ = 0.46, see Table 1 for further statistics). Upon further investigation, we found significant differences between control and stressed rats early on (Day 2: t(29) = 6.51, p < 0.001, d = 2.36), in the middle (Day 6: t(29) = 10.74, p < 0.001, d = 3.88), and at the end (Day 10: t(29) = 10.49, p < 0.001, d = 3.79) of the 10-day stress induction (for results including sex as a between factor, see Table S1 in the Supplement).

**Figure 2.**
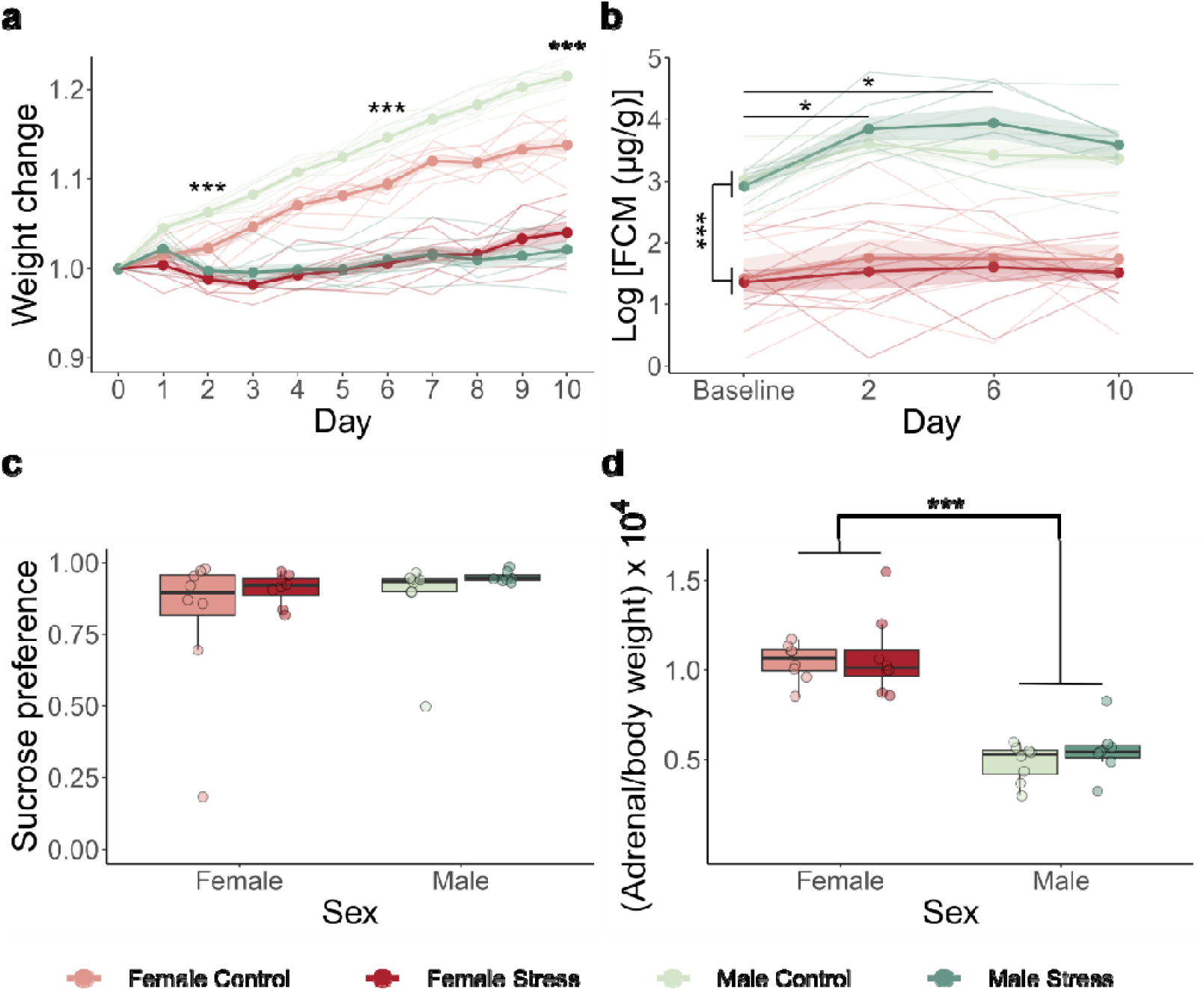
Stress manipulation checks for Experiment 1. The bold lines represent the mean, the surrounding shades are the standard error of the mean and thinner lines represent individual results. **a**. A significant interaction between day and group (F(3.55, 102.87) = 73.42, p < 0.001) in body weight change was detected. There were significant differences in body weight change between the groups starting on day 2 (t(29) = 6.51, p < 0.001). **b**. Males showed higher concentrations of fecal corticosterone metabolites (FCMs) than females (F(1,26) = 146.53, p < 0.001). No group differences were detected (F(1, 26) = 0.16, p = 0.696). **c**. Both groups and both sexes showed a similar preference for the sucrose solution. **d**. Females had a higher adrenal-to-body-weight ratio than males (F(1, 27) = 94.63, p < 0.001), but no differences were found between the control and stress group.

**Table 1.** Stress induction outcomes – Experiment 1.

| Measured outcome | Statistical test | Result | Effect size |
| --- | --- | --- | --- |
| Weight gain | Mixed ANOVA (Greenhouse-Geisser sphericity correction) | Group: $F(1, 29) = 149.98$, $p < 0.001$<br>Day: $F(3.55, 102.87) = 139.05$, $p < 0.001$<br>G*D: $F(3.55, 102.87) = 73.42$, $p < 0.001$ | $\omega_p^2 = 0.83$ [0.72, 1]<br>$\omega_p^2 = 0.62$ [0.56, 1]<br>$\omega_p^2 = 0.46$ [0.39, 1] |
| Weight gain | Simple effects by day: two-sample t-test with Bonferroni correction | Day 2 – Group: $t(29) = 6.51$, $p < 0.001$<br>Day 6 – Group: $t(29) = 10.74$, $p < 0.001$<br>Day 10 – Group: $t(29) = 10.49$, $p < 0.001$ | $d = 2.36$ [1.52, 4.19]<br>$d = 3.88$ [2.95, 5.87]<br>$d = 3.79$ [2.97, 5.68] |
| Sucrose preference ratio | Two-way parametric ANOVA | Sex: $F(1, 27) = 1.02$, $p = 0.321$<br>Group: $F(1, 27) = 2.38$, $p = 0.134$<br>S*G: $F(1, 27) = 0.07$, $p = 0.788$ | $\omega_p^2 = 0$ [0, 1]<br>$\omega_p^2 = 0.04$ [0, 1]<br>$\omega_p^2 = 0$ [0, 1] |
| Adrenal gland-to-body weight ratio | Two-way parametric ANOVA | Sex: $F(1, 27) = 94.63$, $p < 0.001$<br>Group: $F(1, 27) = 0.81$, $p = 0.376$<br>S*G: $F(1, 27) = 0.11$, $p = 0.74$ | $\omega_p^2 = 0.75$ [0.6, 1]<br>$\omega_p^2 = 0$ [0, 1]<br>$\omega_p^2 = 0$ [0, 1] |
| Logarithm of the concentration of fecal corticosterone metabolites | Mixed ANOVA | Sex: $F(1, 26) = 146.53$, $p < 0.001$<br>Group: $F(1, 26) = 0.16$, $p = 0.696$<br>Day: $F(3, 78) = 6.09$, $p < 0.001$<br>S*G: $F(1, 26) = 2.03$, $p = 0.166$<br>S*D: $F(3, 78) = 1.42$, $p = 0.243$<br>G*D: $F(3, 78) = 0.44$, $p = 0.726$<br>S*G*D: $F(3, 78) = 0.57$, $p = 0.6637$ | $\omega_p^2 = 0.84$ [0.73, 1]<br>$\omega_p^2 = 0$ [0, 1]<br>$\omega_p^2 = 0.09$ [0, 1]<br>$\omega_p^2 = 0.04$ [0, 1]<br>$\omega_p^2 = 0$ [0, 1]<br>$\omega_p^2 = 0$ [0, 1]<br>$\omega_p^2 = 0$ [0, 1] |
| Logarithm of the concentration of fecal corticosterone metabolites | Pairwise comparisons with Tukey correction | Day baseline-2: $t(107) = -3.16$, $p = 0.01$<br>Day baseline-6: $t(107) = -3.13$, $p = 0.01$<br>Day baseline-10: $t(107) = -2.30$, $p = 0.104$<br>Day 2-6: $t(107) = 0.03$, $p = 1$<br>Day 2-10: $t(107) = 0.83$, $p = 0.841$<br>Day 6-10: $t(107) = 0.8$, $p = 0.853$ | $d = -0.80$ [-1.32, -0.29]<br>$d = -0.80$ [-1.31, -0.28]<br>$d = -0.59$ [-1.11, -0.08]<br>$d = 0.01$ [-0.5, 0.51]<br>$d = 0.21$ [-0.3, 0.72]<br>$d = 0.21$ [-0.3, 0.72] |

Fecal samples were collected at baseline and at day 2, 6 and 10 of the stress induction or control treatment. The concentration of FCMs was analyzed in each sample (Figure 2b). A strong sex effect was found, with female rats having significantly lower FCM concentrations than males (F(1,26) = 146.53, p < 0.001, ω ^2^ = 0.84). We also found a significant main effect of day (F(3, 78) = 6.09, p < 0.001, ω ^2^ = 0.09), where post-hoc testing revealed a significant increase in FCM concentration between baseline and day 2 (t(107) = −3.16, p = 0.01, d = −0.8) and between baseline and day 6 (t(107) = −3.13, p = 0.01, d = −0.8). We did not find any differences between groups (F(1, 26) = 0.16, p = 0.696, ω ^2^ = 0).

One day after the last session of stress induction or control treatment, we conducted a sucrose preference test (Figure 2c). Chronic restraint stress did not have an effect on the sucrose preference ratio (F(1,27) = 2.38, p = 0.134, ω_p_^2^ = 0.04).

At the end of all experimental manipulations, we dissected the adrenal glands to assess the adrenal gland-to-body weight ratio (Figure 2d). While we did not find any significant differences between the stress and control groups (F(1, 27) = 0.81, p = 0.376, ω_p_^2^ = 0), we did observe a strong sex dimorphism (F(1, 27) = 94.63, p < 0.001, ω_p_^2^ = 0.75), where regardless of group, females had a higher adrenal-to-body-weight ratio than males.

#### Chronically stressed rats show an impairment in acquiring avoidance in the two-way active avoidance procedure, but not in its extinction

To evaluate whether chronic restraint stress has an effect on active avoidance behavior, we subjected rats that underwent chronic restraint and control rats to the two-way active avoidance task. During avoidance acquisition in context A, stressed rats showed a significantly lower number of avoidance responses compared to control rats (F(1, 27) = 9.79, p = 0.004, ω_p_^2^ = 0.23, see Table 2 for further statistics). A significant main effect of day showed that all rats learned the avoidance response over the days (F(2, 54) = 29.31, p < 0.001, ω_p_^2^ = 0.23, Figure 3a). When looking at the number of avoidance responses during the 3 blocks of day 1, main effects of group (F(1, 27) = 5.39, p = 0.028, ω_p_^2^ = 0.13) and block (F(2, 54) = 11.67, p < 0.001, ω_p_^2^ = 0.13) were found (Figure 3b). Rats in the stress group overall thus showed less avoidance responses than rats in the control group on day 1 across all blocks.

**Figure 3.**
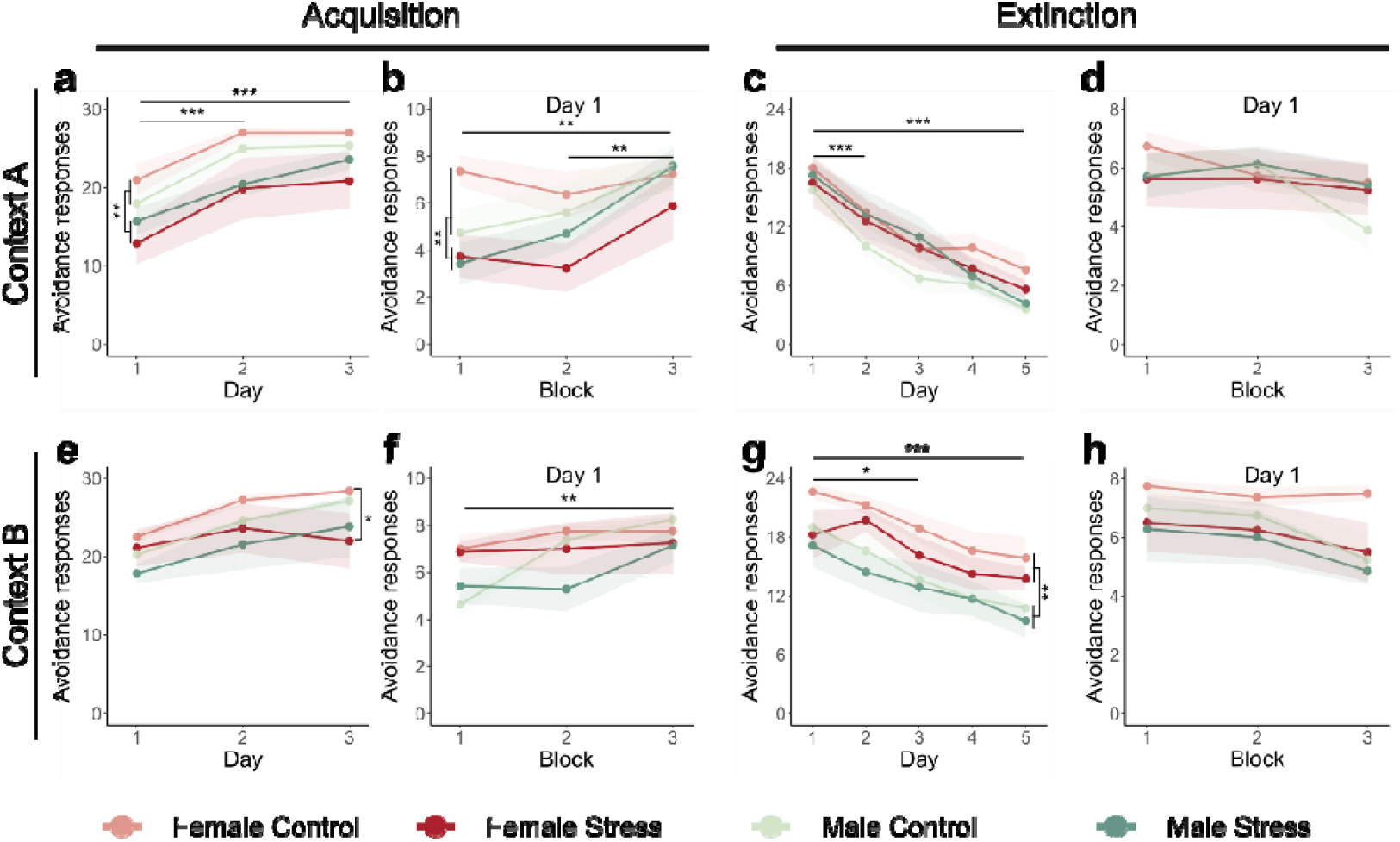
Avoidance responses during two-way active avoidance acquisition and extinction across two distinct contexts in Experiment 1. The lines represent the mean number of avoidance responses, the surrounding shaded area represents the standard error of the mean. Results are expressed in the absolute number of avoidance responses. Each acquisition day consisted of 30 trials, and each extinction day consisted of 24 trials. **a**. Avoidance acquisition in context A. Rats that were subjected to chronic restraint stress showed significantly fewer avoidance responses than rats in the control group throughout all days of acquisition (F(1, 27) = 9.79, p = 0.004). **b**. Results of day 1 avoidance acquisition in context A indicate that rats in the stress group overall showed less avoidance responses than rats in the control group across all blocks (F(1, 27) = 5.39, p = 0.028) **c**. Extinction in context A. All rats, regardless of group or sex, demonstrated a similar rate of extinction of avoidance responses (F(2.79, 75.22) = 55.67, p < 0.001). **d**. Analysis on block level during day 1 of extinction in context A did not reveal any group or sex differences. **e**. Avoidance acquisition in context B. Female rats in the stress group displayed a significantly lower number of avoidance responses compared to female controls on day 3 of acquisition in context B (W = 57.5, p = 0.021). **f**. Analysis on block level during day 1 of acquisition in context B revealed a significant main effect of block (F(2, 54) = 6.01, p = 0.004) meaning that regardless of sex or group, rats learned to acquire avoidance responses. **g**. Extinction in context B. Female rats, regardless of group, showed a higher number of avoidance responses than male rats across all days of extinction in context B (F(1, 27) = 7.34, p = 0.012). **h**. Analysis on block level during day 1 of extinction in context B revealed a significant effect of block, but only in male rats (χ^2^(2)□=□12.61, p = 0.002).

**Table 2.** Avoidance responses during acquisition and extinction in contexts A and B – Experiment 1.

| Test phase | Statistical test | Result | Effect size |
| --- | --- | --- | --- |
| Acquisition context A | Non-parametric mixed ANOVA | Sex: $F(1, 27) = 0.04$, $p = 0.846$<br>Group: $F(1, 27) = 9.79$, **$p = 0.004$**<br>Day: $F(2, 54) = 29.31$, **$p < 0.001$**<br>$S^*G$: $F(1, 27) = 0.79$, $p = 0.382$<br>$S^*D$: $F(2, 54) = 0.40$, $p = 0.67$<br>$G^*D$: $F(2, 54) = 1.32$, $p = 0.277$<br>$S^*G^*D$: $F(2, 54) = 1.05$, $p = 0.355$ | $\omega_p^2 = 0$ [0, 1]<br>$\omega_p^2 = 0.23$ [0.04, 1]<br>$\omega_p^2 = 0.23$ [0.07, 1]<br>$\omega_p^2 = 0.02$ [0, 1]<br>$\omega_p^2 = 0$ [0, 1]<br>$\omega_p^2 = 0$ [0, 1]<br>$\omega_p^2 = 0$ [0, 1] |
| Acquisition context A | Pairwise comparisons with Tukey correction | Day 1-2: $t(54) = -5.74$, **$p < 0.001$**<br>Day 1-3: $t(54) = -7.23$, **$p < 0.001$**<br>Day 2-3: $t(54) = -1.49$, $p = 0.304$ | $r = -0.57$ [-0.74, -0.35]<br>$r = -0.71$ [-0.82, -0.53]<br>$r = -0.07$ [-0.35, 0.21] |
| Acquisition context A day 1 | Mixed ANOVA | Sex: $F(1, 27) = 0$, $p = 0.972$<br>Group: $F(1, 27) = 5.39$, **$p = 0.028$**<br>Block: $F(2, 54) = 11.67$, **$p < 0.001$**<br>$S^*G$: $F(1, 27) = 1.70$, $p = 0.204$<br>$S^*B$: $F(2, 54) = 3.11$, $p = 0.053$<br>$G^*B$: $F(2, 54) = 1.54$, $p = 0.224$<br>$S^*G^*B$: $F(2, 54) = 0.14$, $p = 0.873$ | $\omega_p^2 = 0$ [0, 1]<br>$\omega_p^2 = 0.13$ [0, 1]<br>$\omega_p^2 = 0.13$ [0.01, 1]<br>$\omega_p^2 = 0.02$ [0, 1]<br>$\omega_p^2 = 0.03$ [0, 1]<br>$\omega_p^2 = 0$ [0, 1]<br>$\omega_p^2 = 0$ [0, 1] |
| Acquisition context A day 1 | Pairwise comparisons with Tukey correction | Block 1-2: $t(81) = -0.24$, $p = 0.968$<br>Block 1-3: $t(81) = -3.33$, **$p = 0.004$**<br>Block 2-3: $t(81) = -3.10$, **$p = 0.008$** | $d = -0.06$ [-0.57, 0.44]<br>$d = -0.85$ [-1.37, -0.32]<br>$d = -0.78$ [-1.30, -0.26] |
| Extinction context A | Mixed ANOVA (Greenhouse-Geisser sphericity correction) | Sex: $F(1, 27) = 2.06$, $p = 0.163$<br>Group: $F(1, 27) = 0.13$, $p = 0.719$<br>Day: $F(2.79, 75.22) = 55.67$, **$p < 0.001$**<br>$S^*G$: $F(1, 27) = 2.24$, $p = 0.146$<br>$S^*D$: $F(2.79, 75.22) = 0.51$, $p = 0.664$<br>$G^*D$: $F(2.79, 75.22) = 1.09$, $p = 0.355$<br>$S^*G^*D$: $F(2.79, 75.22) = 0.09$, $p = 0.956$ | $\omega_p^2 = 0.04$ [0, 1]<br>$\omega_p^2 = 0$ [0, 1]<br>$\omega_p^2 = 0.48$ [0.36, 1]<br>$\omega_p^2 = 0.04$ [0, 1]<br>$\omega_p^2 = 0$ [0, 1]<br>$\omega_p^2 = 0$ [0, 1]<br>$\omega_p^2 = 0$ [0, 1] |
| Extinction context A | Pairwise comparisons with Tukey correction | Day 1-2: $t(135) = 4.11$, **$p < 0.001$**<br>Day 1-3: $t(135) = 6.84$, **$p < 0.001$**<br>Day 1-4: $t(135) = 8.35$, **$p < 0.001$**<br>Day 1-5: $t(135) = 10.56$, **$p < 0.001$**<br>Day 2-3: $t(135) = 2.73$, $p = 0.055$ | $d = 1.05$ [0.53, 1.57]<br>$d = 1.74$ [1.20, 2.29]<br>$d = 2.12$ [1.56, 2.69]<br>$d = 2.69$ [2.09, 3.28]<br>$d = 0.70$ [0.19, 1.21] |
| | | Day 2-4: $t(135) = 4.23$, $p < 0.001$<br>Day 2-5: $t(135) = 6.44$, $p < 0.001$<br>Day 3-4: $t(135) = 1.50$, $p = 0.562$<br>Day 3- 5: $t(135) = 3.71$, $p < 0.01$<br>Day 4- 5: $t(135) = 2.21$, $p = 0.183$ | $d = 1.08 [0.56, 1.60]$<br>$d = 1.64 [1.10, 2.18]$<br>$d = 0.38 [-0.12, 0.89]$<br>$d = 0.94 [0.43, 1.46]$<br>$d = 0.56 [0.05, 1.07]$ |
| Extinction context A day 1 | Mixed ANOVA (Greenhouse-Geisser sphericity correction) | Sex: $F(1, 27) = 0.20$, $p = 0.660$<br>Group: $F(1, 27) = 0$, $p = 0.991$<br>Block: $F(1.6, 43.17) = 3.40$, $p = 0.052$<br>S*G: $F(1, 27) = 0.85$, $p = 0.365$<br>S*B: $F(1.6, 43.17) = 1.13$, $p = 0.323$<br>G*B: $F(1.6, 43.17) = 1.15$, $p = 0.317$<br>S*G*B: $F(1.6, 43.17) = 0.52$, $p = 0.558$ | $\omega_p^2 = 0 [0, 1]$<br>$\omega_p^2 = 0 [0, 1]$<br>$\omega_p^2 = 0.04 [0, 1]$<br>$\omega_p^2 = 0 [0, 1]$<br>$\omega_p^2 = 0 [0, 1]$<br>$\omega_p^2 = 0 [0, 1]$<br>$\omega_p^2 = 0 [0, 1]$ |
| Acquisition context B | Non-parametric mixed ANOVA | Sex: $F(1, 27) = 4.48$, $p = 0.044$<br>Group: $F(1, 27) = 0.12$, $p = 0.732$<br>Day: $F(2, 54) = 73.10$, $p < 0.001$<br>S*G: $F(1, 27) = 0.07$, $p = 0.789$<br>S*D: $F(2, 54) = 11.13$, $p < 0.001$<br>G*D: $F(2, 54) = 1.05$, $p = 0.356$<br>S*G*D: $F(2, 54) = 5.95$, $p = 0.005$ | $\omega_p^2 = 0 [0, 1]$<br>$\omega_p^2 = 0.06 [0, 1]$<br>$\omega_p^2 = 0.12 [0.01, 1]$<br>$\omega_p^2 = 0 [0, 1]$<br>$\omega_p^2 = 0 [0, 1]$<br>$\omega_p^2 = 0 [0, 1]$<br>$\omega_p^2 = 0 [0, 1]$ |
| Acquisition context B | Effects by sex (females) - non-parametric mixed ANOVA | Group: $F(1, 14) = 0.22$, $p = 0.647$<br>Day: $F(2, 28) = 30.74$, $p < 0.001$<br>G*D: $F(2, 28) = 11.57$, $p < 0.001$ | $\omega_p^2 = 0.04 [0, 1]$<br>$\omega_p^2 = 0.07 [0, 1]$<br>$\omega_p^2 = 0.02 [0, 1]$ |
| Acquisition context B | Simple effects by day (females): Wilcoxon rank sum test with Bonferroni correction | Day 1 – Group: $W = 26.5$, $p = 1$<br>Day 2 – Group: $W = 33$, $p = 1$<br>Day 3 – Group: $W = 57.5$, $p = 0.021$ | $r = 0.15 [0, 0.62]$<br>$r = 0.03 [0, 0.6]$<br>$r = 0.69 [0.4, 0.86]$ |
| Acquisition context B | Effects by sex (males) - non-parametric mixed ANOVA | Group: $F(1, 13) = 0.27$, $p = 0.614$<br>Day: $F(2, 26) = 45.08$, $p < 0.001$<br>G*D: $F(2, 26) = 1.26$, $p = 0.3$ | $\omega_p^2 = 0.03 [0, 1]$<br>$\omega_p^2 = 0.19 [0, 1]$<br>$\omega_p^2 = 0 [0, 1]$ |
| Acquisition context B | Pairwise comparisons with Tukey correction (males) | Day 1-2: $t(26) = -6.1$, $p < 0.001$<br>Day 1-3: $t(26) = -9.42$, $p < 0.001$<br>Day 2-3: $t(26) = -3.32$, $p = 0.007$ | $r = -0.8 [-0.91, -0.6]$<br>$r = -0.87 [-0.94, -0.72]$<br>$r = -0.67 [-0.84, -0.37]$ |
| Acquisition context B day 1 | Mixed ANOVA | Sex: $F(1, 27) = 3.92$, $p = 0.058$<br>Group: $F(1, 27) = 1.83$, $p = 0.188$<br>Block: $F(2, 54) = 6.01$, $p = 0.004$<br>S*G: $F(1, 27) = 0.13$, $p = 0.718$ | $\omega_p^2 = 0.09 [0, 1]$<br>$\omega_p^2 = 0.03 [0, 1]$<br>$\omega_p^2 = 0.09 [0, 1]$<br>$\omega_p^2 = 0 [0, 1]$ |
| | | S*B: $F(2, 54) = 2.58, p = 0.085$<br>G*B: $F(2, 54) = 1.83, p = 0.170$<br>S*G*B: $F(2, 54) = 0.77, p = 0.468$ | $\omega_p^2 = 0.03 [0, 1]$<br>$\omega_p^2 = 0.02 [0, 1]$<br>$\omega_p^2 = 0 [0, 1]$ |
| Acquisition context B day 1 | Pairwise comparisons with Tukey correction | Block 1-2: $t(81) = -1.73, p = 0.2$<br>Block 1-3: $t(81) = -3.21, p = \mathbf{0.005}$<br>Block 2-3: $t(81) = -1.48, p = 0.305$ | $d = -0.44 [-0.95, 0.07]$<br>$d = -0.82 [-1.34, -0.3]$<br>$d = -0.38 [-0.89, 0.13]$ |
| Extinction context B | Mixed ANOVA | Sex: $F(1, 27) = 7.34, p = \mathbf{0.012}$<br>Group: $F(1, 27) = 1.71, p = 0.203$<br>Day: $F(4, 108) = 24.56, p < \mathbf{0.001}$<br>S*G: $F(1, 27) = 0.22, p = 0.642$<br>S*D: $F(4, 108) = 0.80, p = 0.53$<br>G*D: $F(4, 108) = 0.37, p = 0.083$<br>S*G*D: $F(4, 108) = 0.33, p = 0.854$ | $\omega_p^2 = 0.18 [0.02, 1]$<br>$\omega_p^2 = 0.02 [0, 1]$<br>$\omega_p^2 = 0.22 [0.09, 1]$<br>$\omega_p^2 = 0 [0, 1]$<br>$\omega_p^2 = 0 [0, 1]$<br>$\omega_p^2 = 0 [0, 1]$<br>$\omega_p^2 = 0 [0, 1]$ |
| Extinction context B | Pairwise comparisons with Tukey correction | Day 1-2: $t(135) = 0.97, p = 0.868$<br>Day 1-3: $t(135) = 3.04, p = \mathbf{0.023}$<br>Day 1-4: $t(135) = 4.44, p < \mathbf{0.001}$<br>Day 1-5: $t(135) = 5.33, p < \mathbf{0.001}$<br>Day 2-3: $t(135) = 2.07, p = 0.24$<br>Day 2-4: $t(135) = 3.47, p = \mathbf{0.006}$<br>Day 2-5: $t(135) = 4.35, p < \mathbf{0.001}$<br>Day 3-4: $t(135) = 1.4, p = 0.630$<br>Day 3-5: $t(135) = 2.29, p = 0.156$<br>Day 4-5: $t(135) = 0.89, p = 0.901$ | $d = 0.25 [-0.26, 0.75]$<br>$d = 0.77 [0.26, 1.28]$<br>$d = 1.13 [0.61, 1.65]$<br>$d = 1.36 [0.83, 1.88]$<br>$d = 0.53 [0.02, 1.03]$<br>$d = 0.88 [0.37, 1.4]$<br>$d = 1.11 [0.59, 1.63]$<br>$d = 0.36 [-0.15, 0.86]$<br>$d = 0.58 [0.07, 1.09]$<br>$d = 0.23 [-0.28, 0.73]$ |
| Extinction context B day 1 | Non-parametric mixed ANOVA | Sex: $F(1, 27) = 3.66, p = 0.066$<br>Group: $F(1, 27) = 1.55, p = 0.224$<br>Block: $F(2, 54) = 12.50, p < \mathbf{0.001}$<br>S*G: $F(1, 27) = 0.01, p = 0.944$<br>S*B: $F(2, 54) = 3.63, p = \mathbf{0.033}$<br>G*B: $F(2, 54) = 1.03, p = 0.363$<br>S*G*B: $F(2, 54) = 0.06, p = 0.945$ | $\omega_p^2 = 0.02 [0, 1]$<br>$\omega_p^2 = 0.05 [0, 1]$<br>$\omega_p^2 = 0.05 [0, 1]$<br>$\omega_p^2 = 0 [0, 1]$<br>$\omega_p^2 = 0 [0, 1]$<br>$\omega_p^2 = 0 [0, 1]$<br>$\omega_p^2 = 0 [0, 1]$ |
| Extinction context B day 1 | Friedman rank sum test (female) | Block: $\chi^2(2) = 1.15, p = 0.562$ | $W = 0.04 [0, 0.36]$ |
| Extinction context B day 1 | Friedman rank sum test (male) | Block: $\chi^2(2) = 12.61, p = \mathbf{0.002}$ | $W = 0.42 [0.12, 0.86]$ |
| Extinction context B | Pairwise Wilcoxon signed rank | Block 1-2: $V = 31.5, p = 0.825$ | $r = 0.4 [-0.17, 0.51]$ |
| day 1 | test with Bonferroni correction (males) | Block 1-3: $V = 90$, $p = 0.057$<br>Block 2-3: $V = 68$, $p = 0.069$ | $r = 0.71$ [0.29, 0.90]<br>$r = 0.74$ [0.37, 0.91] |

We hypothesized that rats in the stress group would exhibit more freezing compared to control rats when acclimating to a new environment (the 2WAA shuttle box). However, none of the rats showed any freezing during acclimation on day 1 of acquisition in context A. We did an exploratory analysis on freezing during acclimation on day 2 of acquisition in context A, in which we observed similar low freezing levels across groups and sexes (see Table S2 and Figure S1).

To test for possible effects of chronic restraint stress on general locomotion, we exploratively analyzed the number of crossings during the 5-min acclimation period on the first day of avoidance acquisition in context A, i.e., before any shocks were administered. All rats showed an equal number of crossings during this time period (Table S2 and Figure S2a).

Additionally, we exploratively analyzed the number of crossings during the intertrial intervals, over the acquisition days. Analysis revealed a significant main effect of day (F(2,54) = 3.8, p = 0.029, ω_p_^2^ = 0.04), although the post-hoc test did not identify significant pairwise differences. We also observed a borderline significant main effect of group (F(1,27) = 4.21, p = 0.05, ω_p_^2^ = 0.10), where stressed rats made fewer crossings than control rats during the intertrial intervals (Table S2 and Figure S2b).

To further characterize the behavior of the animals, we exploratively analyzed mean avoidance latency and mean escape latency in acquisition context A. The analysis showed no differences between groups and sexes in mean avoidance latency (Table S2 and Figure S3a). Regarding mean escape latency, only a significant main effect of day was found (F(2, 52) = 3.93, p = 0.026, ω ^2^ = 0.02), but no significant differences were found in the pairwise comparisons of the post-hoc analysis (Table S2 and Figure S3b).

After acquisition in context A, rats underwent 5 daily sessions of extinction training in the same context. Rats from both groups and sexes equally extinguished their avoidance responses over the days (F(2.79, 75.22) = 55.67, p < 0.001, ω_p_^2^ = 0.48, Figure 3c). A significant reduction of avoidance responses was already observed from day 1 to day 2 (t(135) = 4.11, p < 0.001, d = 1.05). Analysis on block level during day 1 of extinction did not reveal any group or sex differences (Figure 3d).

#### Sex differences in avoidance acquisition and extinction in a second 2WAA context

In order to determine whether the effects of chronic restraint stress persist in a new 2WAA learning environment, all rats again went through avoidance training, but this time in a new context B and with a different warning signal (Figure 3e). Analysis revealed a significant sex by day by group interaction (F(2, 54) = 5.95, p = 0.005, ω ^2^ = 0). Follow-up analyses showed that males increased their avoidance responses across both groups over test sessions (F(2, 26) = 45.08, p < 0.001, ω_p_^2^ = 0.19), whereas females showed an interaction between group and day (F(2, 28) = 11.57, p < 0.001, ω_p_^2^ = 0.02). Particularly, stressed females showed a significantly lower number of avoidance responses than control females on the last day of avoidance acquisition in context B (W = 57.50, p = 0.021, r = 0.69). Zooming in on the first day of avoidance training in context B, only a significant main effect of block was found (F(2, 54) = 6.01, p = 0.004, ω ^2^ = 0.09, Figure 3f), meaning that regardless of sex or group, rats learned to acquire avoidance responses. Follow-up analysis showed that a significant increase of avoidance responses occurred from block 1 to 3 (t(81) = −3.21, p = 0.005, d = - 0.82).

We exploratively analyzed mean avoidance latency and mean escape latency during acquisition in context B. There were no significant differences between groups and sexes (Table S2 and Figure S4). We did observe a significant decrease of mean avoidance latency (F(2, 54) = 17.98, p < 0.001, ω ^2^ = 0.19) across days, already from day 1 to day 2 (t(81) = 2.68, p = 0.024, d = 0.68).

Analysis of extinction in context B revealed a sex difference (Figure 3g). A significant main effect of sex showed that female rats had an overall higher number of avoidance responses throughout all extinction sessions, compared to male rats (F(1, 27) = 7.34, p = 0.012, ω_p_^2^ = 0.18). A significant main effect of day revealed that rats from both groups and sexes learned to extinguish their avoidance responses (F(4, 108) = 24.56, p < 0.001, ω_p_^2^ = 0.22). A significant reduction of avoidance responses was observed already early on (e.g., from day 1 to day 3 (t(135) =3.04, p = 0.023, d = 0.77)). At the block level on day 1, we found a significant sex by block interaction (F(2, 54) = 3.63, p = 0.033, ω_p_^2^ = 0, Figure 3h). Although the Friedman test indicated a significant main effect of block (χ^2^(2)LJ=LJ12.61, p = 0.002, W = 0.42) in male rats, post-hoc Wilcoxon signed-rank tests did not identify significant pairwise differences.

We explored whether there were differences in avoidance responding between the first acquisition session of context A and B. We found an interaction between Group and Day (F(1, 27) = 5.46, p = 0.027, ω_p_^2^ = 0.01), but post-hoc tests did not indicate significant differences in number of avoidance responses in stress (V = 27, p = 0.064, r = 0.48) versus control rats (V = 36.5, p = 0.551, r = 0.1). Similarly, we explored whether there were differences in avoidance responding between the last extinction session of context A and B.

We found that females showed more avoidance responses despite extinction (F(1, 27) = 12.35, p = 0.002, ω_p_^2^ = 0.24) than males (females: M = 10.4, SE = 1.07; males: M = 7.86, SE = 0.92). We also observed that rats avoided less at the end of extinction in context A (F(1, 27) = 98.69, p < 0.001, ω_p_^2^ = 0.48) than in context B (context A: M = 5.92, SE = 0.67; context B: M = 13.1, SE= 0.82).

### Experiment 2

#### Chronic restraint stress affects body weight and FCMs, but not sucrose preference or relative adrenal weight

In Experiment 2, rats were food-restricted and trained for lever pressing (see Figure 1b and Methods). Subsequently, rats underwent the 10-day stress induction or control treatment. Body weight change was assessed in all rats (Figure 4a). We found a significant interaction between group and day (F(2.56, 76.81) = 62.98, p < 0.001; for further statistics, see Table 3). Upon further investigation, we found significant differences between control and stressed rats at the beginning (Day 2: t(30) = 6.09, p < 0.001, d = 2.15), middle (Day 6: t(30) = 10.76, p < 0.001, d = 3.8) and end (Day 10: t(30) = 10.22, p < 0.001, d = 3.61) of treatment (for results including sex as within factor, see Table S3). Fecal samples were collected at the same time points during the stress induction phase as in Experiment 1 (Figure 4b). We again found a strong sex effect, with females showing lower FCM concentrations than males (F(1, 28) = 242.92, p < 0.001, ω_p_^2^ = 0.89). In addition, we found an interaction between day and group (F(3, 84) = 3.15, p = 0.029, _p_^2^ = 0.04, see Table 2 for further analyses). Concretely, we did not find any differences between groups at baseline (t(112) = 1.12, p = 0.26, d = 0.4), day 2 (t(112) = −0.71, p = 0.479, d = −0.25) and day 6 (t(112) = −0.47, p = 0.637, d = −0.17). However, on day 10, the stress group showed higher concentrations of FCMs than the control group (t(112) = −2.78, p = 0.006, d =−0.98).

**Figure 4.**
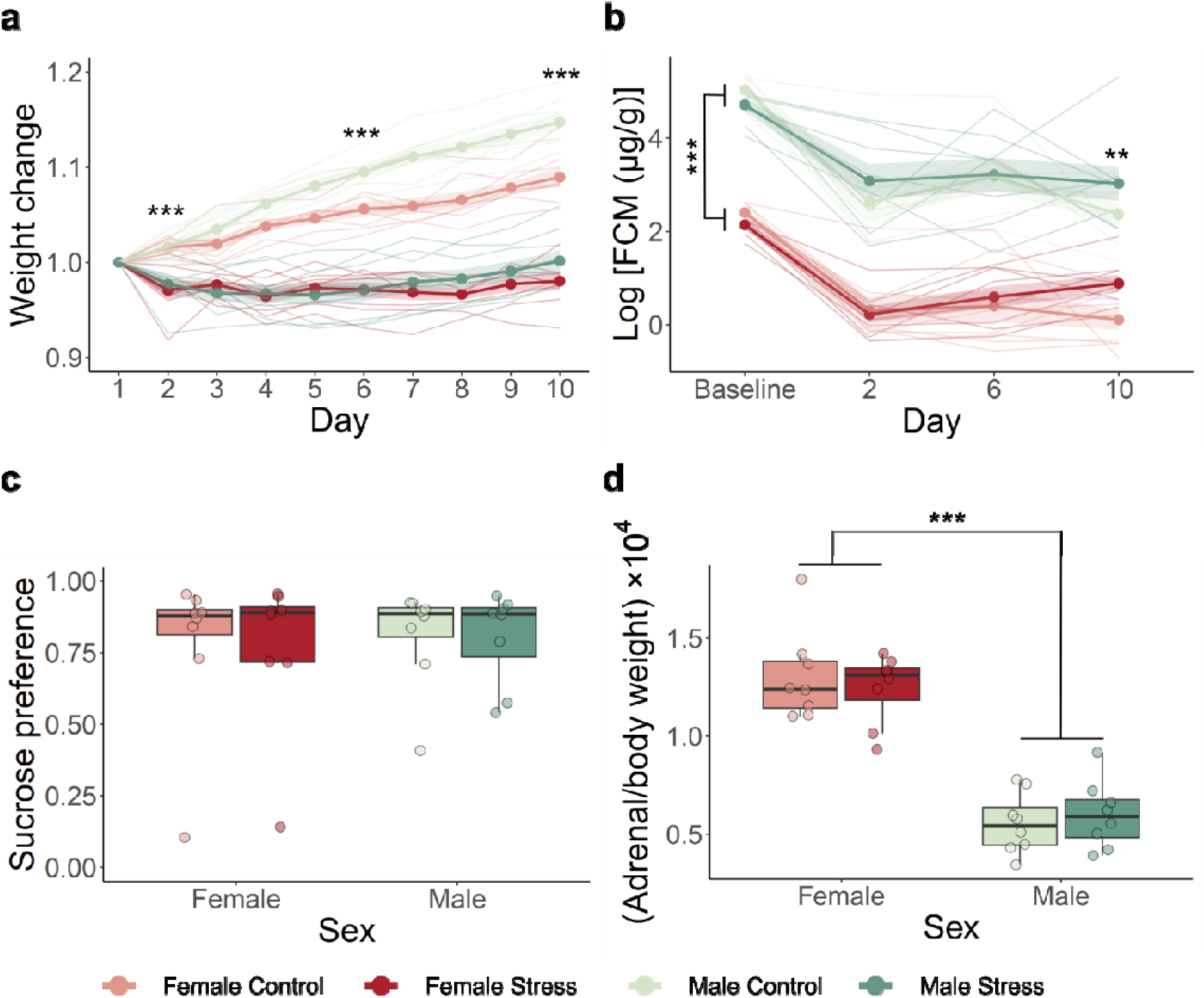
Stress manipulation checks for Experiment 2. The bold lines represent the mean, the surrounding shaded area the standard error of the mean and thinner lines represent individual results. **a.** A significant interaction between day and group (F(2.56, 76.81) = 62.98, p < 0.001) in body weight change was detected. There were significant differences in body weight change between the groups starting on day 2 (t(30) = 6.09, p < 0.001). **b.** Males showed higher concentrations of fecal corticosterone metabolites (FCMs) than females (F(1, 28) = 242.92, p < 0.001). At day 10, stress animals had higher concentrations of FCMs than their control counterparts (t(112) = −2.78, p = 0.006)**. c.** Both groups and both sexes showed a similar preference for the sucrose solution. **d.** Females had higher adrenal gland-to-body weight ratios than male animals (F(1, 27) = 133.45, p < 0.001), but no differences were observed between control and stress animals.

**Table 3.** Stress induction outcomes – Experiment 2.

| Measured outcome | Statistical Test | Result | Effect size |
| --- | --- | --- | --- |
| Weight change | Mixed ANOVA (Greenhouse-Geisser sphericity correction) | Group: $F(1, 30) = 123.64$, $p < 0.001$<br>Day: $F(2.56, 76.81) = 61.1$, $p < 0.001$<br>$G^*D$: $F(2.56, 76.81) = 62.98$, $p < 0.001$ | $\omega_p^2 = 0.79$ [0.67, 1]<br>$\omega_p^2 = 0.36$ [0.27, 1]<br>$\omega_p^2 = 0.37$ [0.28, 1] |
| Weight change | Simple effects by day: two-sample t-test | Day 2 – Group: $t(30) = 6.09$, $p < 0.001$<br>Day 6 – Group: $t(30) = 10.76$, $p < 0.001$<br>Day 10 – Group: $t(30) = 10.22$, $p < 0.001$ | $d = 2.15$ [1.63, 3.24]<br>$d = 3.8$ [3.03, 5.15]<br>$d = 3.61$ [2.1, 5.43] |
| Sucrose preference ratio | Two-way non-parametric ANOVA | Sex: $F(1, 28) < 0.01$, $p = 0.943$<br>Group: $F(1, 28) = 0.01$, $p = 0.91$<br>$S^*G$: $F(1, 28) < 0.01$, $p = 0.942$ | $\omega_p^2 = 0.03$ [0, 1]<br>$\omega_p^2 = 0.02$ [0, 1]<br>$\omega_p^2 = 0.03$ [0, 1] |
| Logarithm of the concentration of fecal corticosterone metabolites | Mixed ANOVA (Greenhouse-Geisser sphericity correction) | Group: $F(1, 28) = 1.29$, $p = 0.265$<br>Sex: $F(1, 28) = 242.92$, $p < 0.001$<br>$G^*S$: $F(1, 28) = 0.03$, $p = 0.868$<br>Day: $F(3, 84) = 67.8$, $p < 0.001$<br>$G^*D$: $F(3, 84) = 3.15$, $p = 0.029$<br>$S^*D$: $F(3, 84) = 0.89$, $p = 0.45$<br>$G^*S^*D$: $F(3, 84) = 0.48$, $p = 0.694$ | $\omega_p^2 < 0.01$ [0, 1]<br>$\omega_p^2 = 0.89$ [0.82, 1]<br>$\omega_p^2 = 0$ [0, 1]<br>$\omega_p^2 = 0.58$ [0.47, 1]<br>$\omega_p^2 = 0.04$ [0, 1]<br>$\omega_p^2 = 0$ [0, 1]<br>$\omega_p^2 = 0$ [0, 1] |
| Logarithm of the concentration of fecal corticosterone metabolites | Simple effects by day: two-sample t-test | Baseline – Group: $t(112) = 1.12$, $p = 0.26$<br>Day 2 – Group: $t(112) = -0.71$, $p = 0.479$<br>Day 6 – Group: $t(112) = -0.47$, $p = 0.637$<br>Day 10 – Group: $t(112) = -2.78$, $p = 0.006$ | $d = 0.4$ [-0.31, 1.1]<br>$d = -0.25$ [-0.95, 0.45]<br>$d = -0.17$ [-0.87, 0.53]<br>$d = -0.98$ [-1.69, -0.27] |
| Adrenal gland-to-body weight ratio | Two-way parametric ANOVA | Sex: $F(1, 28) = 112.67$, $p < 0.001$<br>Group: $F(1, 28) = 0.02$, $p = 0.900$<br>$S^*G$: $F(1, 28) = 0.64$, $p = 0.432$ | $\omega_p^2 = 0.78$ [0.64, 1]<br>$\omega_p^2 = 0$ [0, 1]<br>$\omega_p^2 = 0$ [0, 1] |

One day after the last session of stress induction or control treatment, we conducted a sucrose preference test (Figure 4c). We did not observe a significant group difference (F(1, 28) = 0.01, p = 0.91, ω ^2^ = −0.02), but the stress group presented higher variability than the control group.

At the end of all experimental manipulations, we determined the adrenal gland-to-body weight ratio (Figure 4d). While we did not find any significant differences between the stress and control groups (F(1, 28) = 0.02, p = 0.900, ω_p_^2^ = 0), we did observe a strong sex dimorphism (F(1, 28) = 112.67, p < 0.001, ω_p_^2^ = 0.81), where regardless of group, females had a higher adrenal-to-body weight ratio than males.

#### Subtle effects of chronic restraint stress on platform-mediated avoidance acquisition and extinction

Rats underwent platform-mediated avoidance acquisition for 10 days in both groups (Figure 5, see Figure S5 for trial-by-trial data). On each day, 9 CS-US pairings were presented, and rats had the opportunity to avoid or escape from the foot shock US by stepping onto the platform, at the expense of access to a lever which they could press for food.

**Figure 5.**
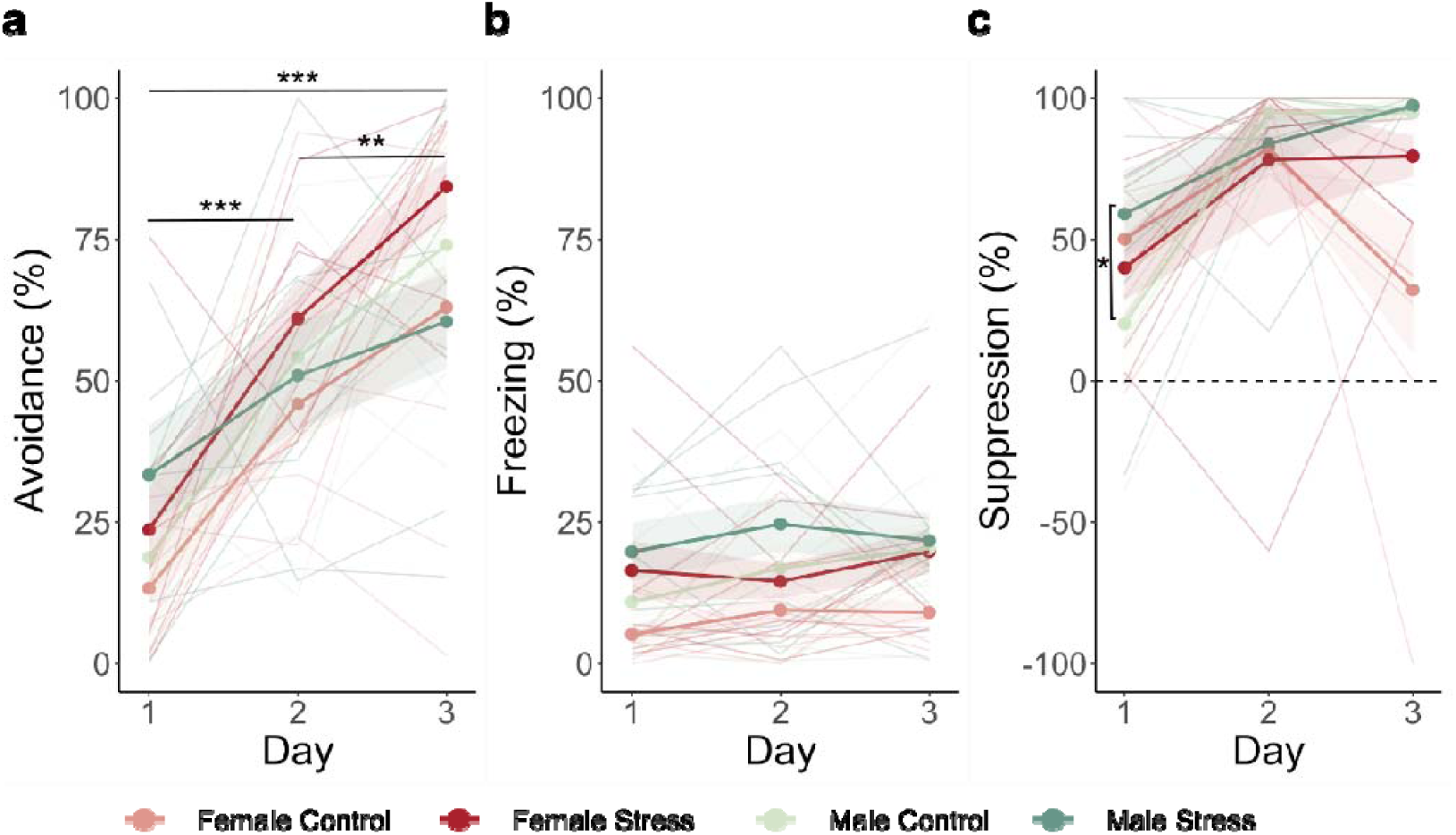
Avoidance acquisition results for Experiment 2. The bold lines represent the mean of the first block (first 3 CS) on each day, the surrounding shaded area the standard error of the mean and thinner lines represent individual results. **a.** Avoidance behavior significantly increased over acquisition sessions (F(2, 56) = 36.73, p < 0.001). **b.** Freezing did not significantly increase over acquisition sessions (F(2, 56) = 1.59, p = 0.212). **c.** Suppression of lever pressing showed a triple interaction of day, sex and group (F(2, 56) = 7.33, p = 0.001). Female rats showed a similar increase of suppression of lever pressing across groups (F(2, 28) = 8.84, p = 0.001), while data for male rats showed an interaction between day and group (F(2, 28) = 10.29, p < 0.001). Stressed males showed significantly higher suppression of lever pressing than control males on day 1 (W = 12, p = 0.04).

We hypothesized that stressed rats would show more avoidance than control rats, but this hypothesis was not confirmed; all rats similarly acquired avoidance responding over the first three acquisition sessions (F(2, 56) = 36.73, p < 0.001, ω_p_^2^ = 0.43; for further statistics, see Table 2). Freezing did not significantly increase over the first three acquisition sessions (F(2, 56) = 1.59, p = 0.212, ω_p_^2^= 0). Regarding suppression of lever pressing, we found a triple interaction between day, sex and group (F(2, 56) = 7.33, p = 0.001, ω_p_^2^ = 0.05). Follow-up analyses showed that females increased their suppression of lever pressing similarly in both groups over the first three test sessions (F(2, 28) = 8.84, p = 0.001, ω ^2^ = 0.08), whereas males showed an interaction between group and day (F(2, 28) = 10.29, p < 0.001, ω_p_^2^ = 0.01). Stressed males showed significantly higher suppression of lever pressing than control males on day 1 (W = 12, p = 0.04, r = −0.62), but no longer on the following days (see Table 4). For exploratory purposes, we also examined time spent on the platform before CS onset, but did not observe clear patterns of behavior distinguishing stressed rats from controls (Figure S6).

**Table 4.**
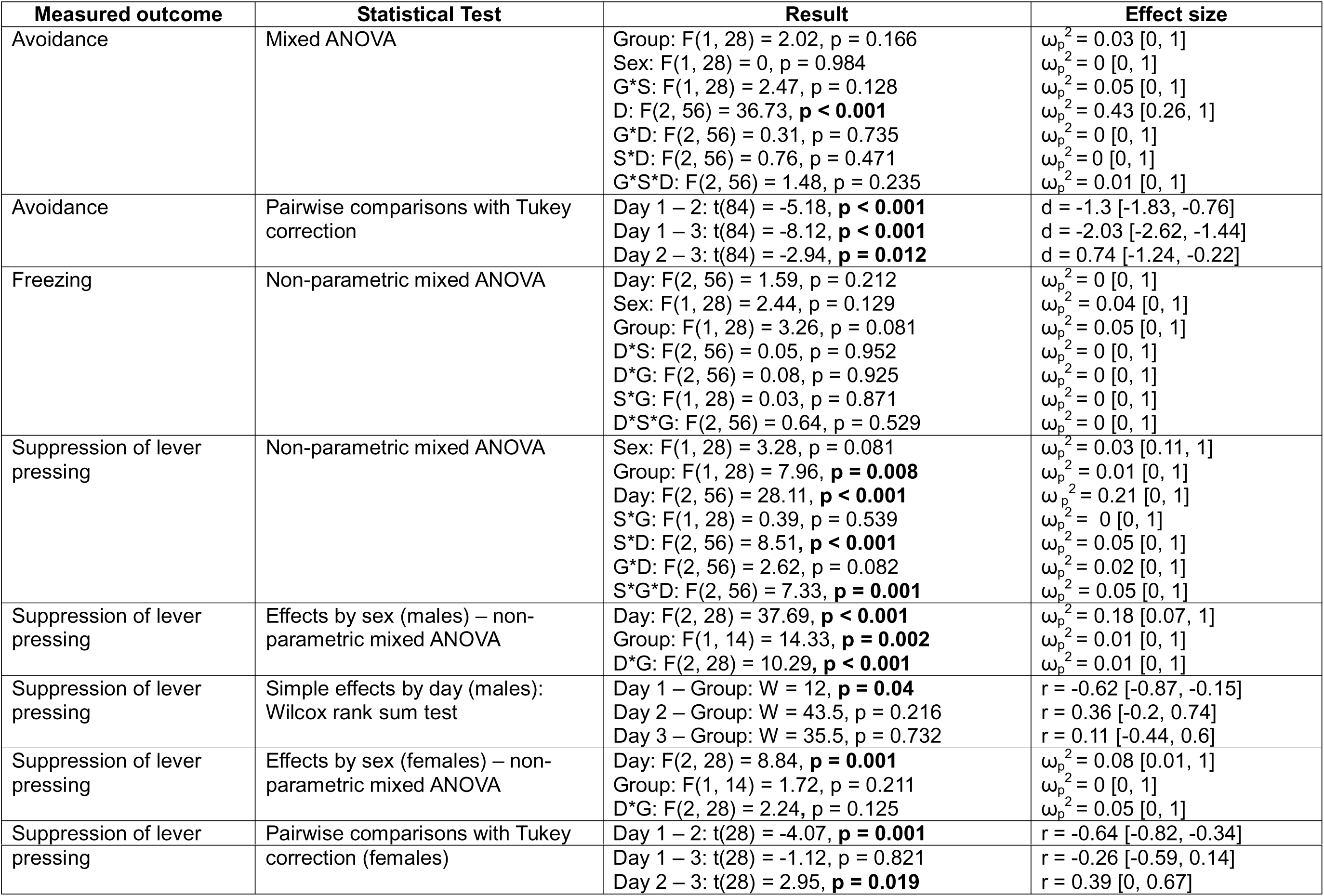
Statistical test results Acquisition – Experiment 2.

We also analyzed darting behavior during the acquisition sessions and found that 37.5% (6/16) of females and 18.75% (3/16) of males were classified as darters (see Figure S7). A Chi-square test indicated that females were not more likely to be classified as darters compared to males (χ^2^(1) = 0.62, p = 0.432). Additionally, we exploratorily investigated whether stress or control rats were more likely to be classified as darters. We found that control rats were more likely to be darters (χ^2^(1) = 5.57, p = 0.018), as only one of the stressed rats (6.25%, or 1/16) was classified as a darter, while half of the control rats (8/16) were classified as darters. Given that in the stress group only one rat was classified as a darter, we only explored in the control group if there were differences between darters and non-darters in defensive behaviors. We did not find any differences in avoidance, freezing and suppression of lever pressing between darters and non-darters during acquisition nor extinction of avoidance (see Table S4 for further statistics).

Next, we performed 4 sessions of extinction training, on consecutive days, with 9 CSs per session (Figure 6, see Figure S8 for trial-by-trial data). We examined how defensive responding to the CS changed over blocks in the first two sessions. Overall, avoidance behavior decreased from the first to the second extinction session (F(1, 140) = 140.88, p < 0.001, ω_p_^2^ = 0.27; for further statistics, see Table 5). We found an interaction between sex and day in freezing (F(1, 140) = 6.53, p = 0.012, ω _p_^2^ = 0.01). Concretely, females showed a significant reduction of freezing over days (V = 705, p < 0.001, r = 0.53), while males did not (V = 644, p = 0.08, r = 0.26). Suppression of lever pressing decreased over the two extinction sessions (F(1, 28) = 26.69, p < 0.001, ω_p_^2^ = 0.12), without significant differences between sexes or groups.

**Figure 6.**
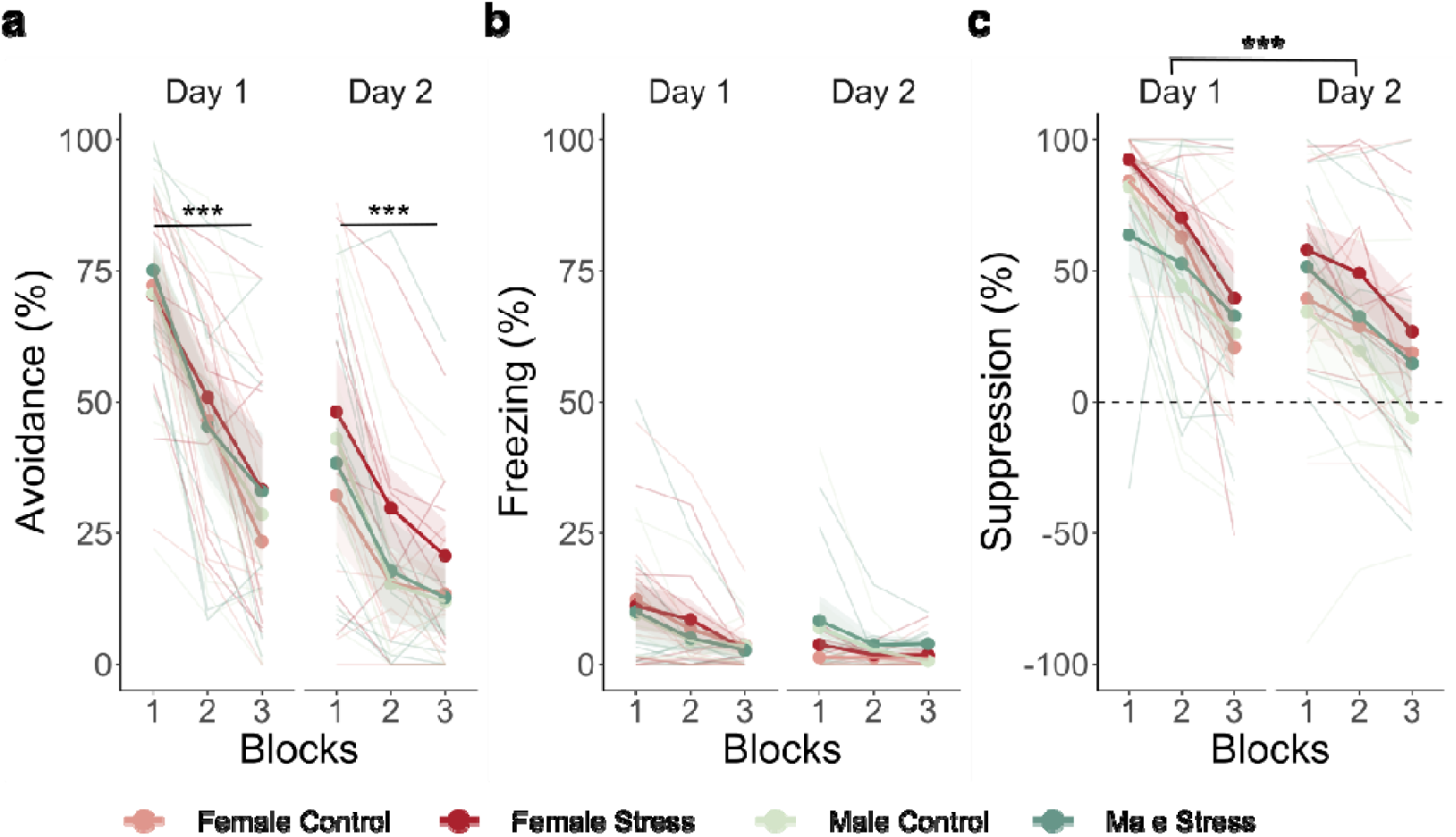
Extinction results for Experiment 2. The bold lines represent the mean of the blocks on each day, the surrounding shaded area the standard error of the mean and thinner lines represent individual results. **a.** Avoidance behavior significantly decreased over the blocks on day 1 (χ^2^(2) = 51.94, p < 0.001) and on day 2 (χ^2^(2) = 37.1, p < 0.001), without any group or sex differences. **b.** Freezing significantly decreased over extinction sessions for females (V = 705, p < 0.001), but not males (V = 644, p = 0.08), in the absence of group differences. **c.** Suppression of lever pressing significantly decreased over extinction sessions (F(1, 28) = 26.69, p < 0.001), without any group or sex differences.

**Table 5.** Statistical test results Extinction – Experiment 2.

| Measured outcome | Statistical Test | Result | Effect size |
| --- | --- | --- | --- |
| Avoidance | Mixed ANOVA | Group: $F(1, 28) = 0.39$, $p = 0.539$<br>Sex: $F(1, 28) = 0.22$, $p = 0.643$<br>Day: $F(1, 140) = 140.88$, **$p < 0.001$**<br>Block: $F(2, 140) = 81.77$, **$p < 0.001$**<br>G*S: $F(1, 28) = 0.29$, $p = 0.593$<br>G*D: $F(1, 140) = 0.4$, $p = 0.528$<br>S*D: $F(1, 140) = 0.7$, $p = 0.403$<br>G*B: $F(2, 140) = 0.01$, $p = 0.988$<br>S*B: $F(2, 140) = 0.74$, $p = 0.48$<br>D*B: $F(2, 140) = 7.34$, **$p < 0.001$**<br>G*S*D: $F(1, 140) = 2.64$, $p = 0.106$<br>G*S*B: $F(2, 140) = 0.12$, $p = 0.89$<br>G*D*B: $F(2, 140) = 0.49$, $p = 0.612$<br>S*D*B: $F(2, 140) = 0.01$, $p = 0.985$<br>G*S*D*B: $F(2, 140) = 0.84$, $p = 0.435$ | $\omega_p^2 = 0$ [0, 1]<br>$\omega_p^2 = 0$ [0, 1]<br>$\omega_p^2 = 0.27$ [0.06, 1]<br>$\omega_p^2 = 0.33$ [0.16, 1]<br>$\omega_p^2 = 0$ [0, 1]<br>$\omega_p^2 = 0$ [0, 1]<br>$\omega_p^2 = 0$ [0, 1]<br>$\omega_p^2 = 0$ [0, 1]<br>$\omega_p^2 = 0$ [0, 1]<br>$\omega_p^2 = 0.02$ [0, 1]<br>$\omega_p^2 < 0.01$ [0, 1]<br>$\omega_p^2 = 0$ [0, 1]<br>$\omega_p^2 = 0$ [0, 1]<br>$\omega_p^2 = 0$ [0, 1]<br>$\omega_p^2 = 0$ [0, 1] |
| Avoidance | Simple effects by block: Day 1 – Friedman test | $X^2(2) = 51.94$, **$p < 0.001$** | $W = 0.81$ [0.66, 1] |
| Avoidance | Simple effects by block: Day 2 – Friedman test | $X^2(2) = 37.1$, **$p < 0.001$** | $W = 0.58$ [0.37, 1] |
| Freezing | Non-parametric mixed ANOVA | Group: $F(1, 28) = 0.48$, $p = 0.49$<br>Sex: $F(1, 28) = 0.26$, $p = 0.614$<br>Day: $F(1, 140) = 15.92$, **$p < 0.001$**<br>Block: $F(2, 140) = 4.85$, **$p = 0.009$**<br>G*S: $F(1, 28) = 0.03$, $p = 0.86$<br>G*D: $F(1, 140) = 0.01$, $p = 0.924$<br>S*D: $F(1, 140) = 6.53$, **$p = 0.012$**<br>G*B: $F(2, 140) = 0.14$, $p = 0.872$<br>S*B: $F(2, 140) = 2.05$, $p = 0.132$<br>D*B: $F(2, 140) = 4.14$, **$p = 0.018$** | $\omega_p^2 = 0$ [0, 1]<br>$\omega_p^2 = 0$ [0, 1]<br>$\omega_p^2 = 0.04$ [0, 1]<br>$\omega_p^2 = 0.06$ [0.01, 1]<br>$\omega_p^2 = 0$ [0, 1]<br>$\omega_p^2 = 0$ [0, 1]<br>$\omega_p^2 = 0.01$ [0, 1]<br>$\omega_p^2 = 0$ [0, 1]<br>$\omega_p^2 = 0$ [0, 1]<br>$\omega_p^2 < 0.01$ [0, 1] |
| | | $G*S*D: F(1, 140) = 0.77, p = 0.381$<br>$G*S*B: F(2, 140) = 0.41, p = 0.661$<br>$G*D*B: F(2, 140) = 0.76, p = 0.468$<br>$S*D*B: F(2, 140) = 0.92, p = 0.4$<br>$G*S*D*B: F(2, 140) = 0.16, p = 0.853$ | $\omega_p^2 = 0 [0, 1]$<br>$\omega_p^2 = 0 [0, 1]$<br>$\omega_p^2 = 0 [0, 1]$<br>$\omega_p^2 = 0 [0, 1]$<br>$\omega_p^2 = 0 [0, 1]$ |
| Freezing | Simple effects by Day: Wilcoxon signed rank test (males) | $V = 644, p = 0.08$ | $r = 0.26 [0.03, 0.53]$ |
| Freezing | Simple effects by Day: Wilcoxon signed rank test (females) | $V = 705, p < 0.001$ | $r = 0.53 [0.29, 0.71]$ |
| Suppression | Mixed ANOVA | $Group: F(1, 28) = 0.87, p = 0.358$<br>$Sex: F(1, 28) = 1.06, p = 0.312$<br>$G*S: F(1, 28) = 0.06, p = 0.811$<br>$Day: F(1, 28) = 26.69, p < 0.001$<br>$G*D: F(1, 28) = 1.28, p = 0.268$<br>$S*D: F(1, 28) = 0.01, p = 0.92$<br>$G*S*D: F(1, 135) = 0.5, p = 0.486$<br>$Block: F(1.63, 45.56) = 41.17, p < 0.001$<br>$G*B: F(1.63, 45.56) = 0.35, p = 0.661$<br>$S*B: F(1.63, 45.56) = 0.24, p = 0.745$<br>$D*B: F(1.63, 45.56) = 0.34, p = 0.672$<br>$G*S*B: F(1.54, 43.25) = 2.06, p = 0.149$<br>$G*D*B: F(1.54, 43.25) = 0.7, p = 0.466$<br>$S*D*B: F(1.54, 43.25) = 1.3, p = 0.276$<br>$G*S*D*B: F(1.54, 43.25) = 0.56, p = 0.533$ | $\omega_p^2 = 0 [0, 1]$<br>$\omega_p^2 < 0.01 [0, 1]$<br>$\omega_p^2 = 0 [0, 1]$<br>$\omega_p^2 = 0.12 [0, 1]$<br>$\omega_p^2 < 0.01 [0, 1]$<br>$\omega_p^2 = 0 [0, 1]$<br>$\omega_p^2 = 0 [0, 1]$<br>$\omega_p^2 = 0.19 [0.05, 1]$<br>$\omega_p^2 = 0 [0, 1]$<br>$\omega_p^2 = 0 [0, 1]$<br>$\omega_p^2 < 0.01 [0, 1]$<br>$\omega_p^2 = 0 [0, 1]$<br>$\omega_p^2 = 0 [0, 1]$<br>$\omega_p^2 < 0.01 [0, 1]$<br>$\omega_p^2 = 0 [0, 1]$ |

In addition, we investigated if there were differences in persistence of avoidance between control and stress groups (see Figure S9). We classified a rat as persistent avoider if it spent more than 50% of the CS duration on the platform, during the first block of the third extinction session. We found that 18.75% of rats (6/32; 3 males and 3 females) were considered persistent avoiders and a Chi-square test (χ^2^ (1) = 0.2, p = 0.651) indicated that neither control nor stressed rats had more chances of being classified as persistent avoiders.

We also investigated whether persistent versus non-persistent avoiders perhaps differed regarding other defensive behaviors. At the end of acquisition (day 10), we did not find any differences in freezing between persistent and non-persistent avoiders (W = 64, p = 0.514, r = 0.012) nor in suppression of lever pressing (W = 104.5, p = 0.202, r = 0.23). However, during extinction, we did observe differences in freezing (W = 22, p = 0.005, r = 0.5) and suppression of lever pressing (W = 3, p < 0.001, r = 0.64), but only on day 3. Concretely, persistent avoiders showed higher freezing (M = 6.75, SE = 2.65) and higher suppression of lever pressing (M = 92.4, SE = 3.86) than non-persistent avoiders (freezing: M = 2.20, SE = 1.11; suppression of lever pressing: M = 7.75, SE = 7.61).

## Discussion

We investigated the effects of chronic restraint stress on the acquisition and extinction of two-way active avoidance (Experiment 1) and platform-mediated avoidance (Experiment 2) in male and female rats. In both experiments, the chronic restraint stress procedure significantly diminished weight gain in stressed rats relative to controls, without affecting sucrose preference or relative adrenal gland weights. A marked sex difference was observed both in relative adrenal gland weight and levels of fecal corticosterone metabolites (FCMs). Female rats had a higher adrenal gland-to-body weight ratio than male rats, as reported previously (Goel et al., 2014). Regarding FCMs, we found that male rats had higher levels than female rats overall, in line with previous literature (Lepschy et al., 2007). Additionally, in Experiment 2, stressed rats showed increased levels of FCMs compared to controls at the end of the stress induction procedure, suggesting that the 10 days of stress induction affected corticosterone levels.

In Experiment 1, previously stressed rats showed an impairment in two-way active avoidance acquisition compared to controls. During subsequent extinction of avoidance, no differences between stressed and control rats were observed. During the next phase of the experiment, which took place in a different context and with a different warning signal, the effects of chronic restraint stress on avoidance acquisition were more limited and sex-specific. Only female stressed rats showed an impairment in avoidance acquisition compared to controls, and only during the last session. During subsequent extinction, females showed overall more avoidance responses than males, regardless of prior exposure to chronic restraint stress.

In contrast with Experiment 1, Experiment 2 showed only subtle effects of chronic stress induction on the acquisition of avoidance. There was no effect on avoidance itself nor on freezing behavior. However, we did find an effect on the suppression of lever pressing, where male stress rats showed higher suppression of lever pressing on day 1 than control males.

Additionally, rats in the stress condition were less likely to be classified as darters. Both males and females readily extinguished avoidance and other defensive behaviors during the extinction phase, although males did not show a significant reduction in freezing over extinction. This lack of a reduction may, however, not be very informative as freezing levels remained rather low throughout the platform-mediated avoidance procedure.

With Experiment 1, we have replicated findings from several authors (Bravo et al., 2009; Dagnino-Subiabre et al., 2005; Gamaro et al., 1999; Ulloa et al., 2010), which support that chronic restraint stress impairs the acquisition of active avoidance behavior in the shuttle avoidance task. This effect has been observed despite differences in chronic restraint stress procedures and, in our case, despite not replicating all stress manipulation checks. Three earlier active avoidance studies reported stress manipulation checks (Gamaro et al. (1999) did not). Dagnino-Subiabre et al. (2005) conducted restraint stress for 2 h daily for 10 days, while Ulloa et al. (2010) and Bravo et al. (2009) reported 2.5 h of daily restraint stress for 13 and 14 days, respectively. The latter studies used plexiglass tubes, presumably similar to our own restraint tubes, and Dagnino-Subiabre et al. (2005) used their own custom immobilization cages. Dagnino-Subiabre et al. (2005) and Ulloa et al. (2010) reported significantly reduced weight gain in stressed rats and an increase in adrenal gland weight in stressed rats compared to controls. Ulloa et al. (2010) examined adrenal glands two days after the last session of chronic stress induction, compared to our 15-16 days after the last stress induction session. In line with our findings, Bravo et al. (2009) observed significant differences in body weight gain but no significant differences in adrenal gland weights between control and stressed rats. Unlike us, Bravo et al. (2009) also found significant differences in a sucrose preference test, with stressed rats showing a reduced preference for the sucrose solution compared to controls. Likewise, a recent meta-analysis indicates that chronic restraint stress decreases sucrose preference in male rodents (Mao et al., 2022). It should be noted that the use of the sucrose preference test as a measure of anhedonia is not uncontested, especially when confounding variables such as metabolic factors are not controlled (Berrio et al., 2024).

It is worth noting that stress induction was conducted during the morning of a standard light-dark cycle in all three studies mentioned above, while half of our rats were restrained in the morning and the other half in the early afternoon. Moreover, they all used male Sprague-Dawley rats, whereas we used male and female Wistar rats. All these differences, alongside the larger sample size in Dagnino-Subiabre et al. (2005) and the longer stress induction procedures in Ulloa et al. (2010) and Bravo et al. (2009), may partially explain why we replicated the effect of chronic restraint stress on weight gain, but not on relative adrenal gland weight or on sucrose preference.

As mentioned above, Experiment 1 replicated the previously described effect of chronic restraint stress impairing acquisition of two-way active avoidance. We added several phases to our procedure to study this effect more extensively. During subsequent acquisition in a novel context B, a significant group difference emerged on the final day of acquisition, with stressed females demonstrating significantly lower avoidance responses than their control counterparts. Albeit more subtle and sex-specific, these findings do suggest that in the second context, chronic stress continued to impact avoidance learning in female rats. During extinction in context B (unlike in context A), female rats—regardless of group—displayed impaired extinction compared to male rats (in line with (Beatty et al., 1971; Landin and Chandler, 2023; Shanazz et al., 2022). When exploratively comparing the number of avoidance responses between end of extinction (day 5) in context A and end of extinction in context B, we observed that rats still avoided more in context B than in context A, and females more than males. Training and extinction in a new context B with the same aversive stimulus as before (foot shock) following initial extinction in context A may have created additional ambiguity and led to reduced extinction efficacy in context B, reflecting a “better-safe-than-sorry” strategy (Van den Bergh et al., 2021).

The lack of effects of chronic restraint stress on the platform-mediated avoidance task in Experiment 2 is surprising. We hypothesized that chronically stressed rats would show increased avoidance in this task. Previous research in humans found that acute stress increased costly avoidance in healthy humans (Vogel and Schwabe, 2019) and that patients suffering from PTSD were biased towards avoidance in an approach-avoidance conflict task compared to controls (Weaver et al., 2020). However, we did not observe significant differences in the acquisition of avoidance behavior between stressed and control rats. An open question for future research is whether such differences might emerge in a variation of the platform-mediated avoidance task that involves a stronger conflict, where food rewards are only available during CS presentation (Bravo-Rivera et al., 2021). In addition, it should be noted that food restriction in itself (Chacón et al., 2005; Heiderstadt et al., 2000) may constitute a prior and continued chronic stressor in the platform-mediated avoidance task, which may mask the effects of purposeful stress induction manipulations. This notion may be supported by Figure 4b, which shows a decrease in fecal corticosterone metabolite levels from baseline (when rats are food-restricted) to day 2 of stress or control treatment (when rats are not food-restricted). Thus, food restriction might have an important effect on corticosterone levels, although this should be examined in more detail, and taking into consideration metabolic differences between food-restricted and free-fed animals. In any case, further studies investigating the effects of stress on platform-mediated avoidance may want to take these reflections into account.

Because of the inherent conflict to approach food and avoid shocks, avoidance in the platform-mediated avoidance task comes at a relatively high cost compared to avoidance in shuttle avoidance tasks. Prior evidence suggests that acute stress exacerbates costly avoidance in individuals with anxiety (Vogel and Schwabe, 2019), but, until now, the effects of stress on costly avoidance in rodents had not yet been investigated. Previous research with rodents did show that chronic stress can lead to reduced exploration of the open arms in the elevated plus maze, which is a low-cost approach-avoidance conflict task (Bondi et al., 2008; Chiba et al., 2012), although other studies did not find such effects (Cox et al., 2011; Mitra et al., 2005).

In addition to the difference in cost between two-way active avoidance and platform-mediated avoidance, there is an additional crucial difference. In the former, there is no safe area, whereas in the latter, the platform is always safe. Additionally, shuttling during the tone in the two-way active avoidance task typically terminates the CS tone, which is not the case in the platform-mediated avoidance task (Diehl et al., 2019). Previous research has shown that stress controllability may recruit a different neurocircuitry than safety learning (Christianson et al., 2008). Shuttle avoidance may primarily tap into rats’ ability to control their environment, without affording strong safety learning, while platform-mediated avoidance may primarily tap into safety learning structures. This divergence could provide an alternative explanation, next to how costly avoidance responses are, for the differential effects of chronic restraint stress on these two active avoidance tasks.

To conclude, chronic restraint stress resulted in acute effects on weight gain in both experiments. In subsequent behavioral testing, prior stress was found to impair avoidance acquisition in the two-active avoidance task, but not in the platform-mediated avoidance task. Extinction of avoidance was not impaired by prior stress in any of the tasks.

## Acknowledgements

We thank Markus Wöhr for providing the USV equipment. Graphical representations in the figures were created with BioRender.com.

## Author contributions

ALM: conceptualization, data curation, formal analysis, investigation, methodology, project administration, software, validation, visualization, writing – original draft. MDC: data curation, formal analysis, investigation, methodology, project administration, software, visualization, writing – original draft. YVH: data curation, investigation; writing – review & editing. LV: methodology, software, writing – review & editing. RP: investigation, methodology, writing – review & editing. BV: funding acquisition, resources, supervision, writing – review & editing. TB: conceptualization, funding acquisition, methodology, project administration, resources, supervision, writing – review & editing. LL: conceptualization, funding acquisition, methodology, project administration, resources, supervision, writing – review & editing. ALM and MDC are joint first authors.

## Funding sources

This work was supported by KU Leuven Research Grant 3H190245 (awarded to TB, LL, BV), FWO PhD fellowship 11K3823N (awarded to ALM), and FWO PhD fellowship 11G7122N (awarded to LV).

## Declaration of competing interests

The authors declare no other conflicts of interest.

## Notes

https://osf.io/qm6y7/

### Competing Interest Statement

The authors have declared no competing interest.

### Summary of Updates

This revision contains new analyses, supplementary materials and an updated introduction and discussion.

