## Supplementary information for "Differential effects of chronic restraint stress on two active avoidance tasks in rats"

Supplementary Figures
Experiment 1

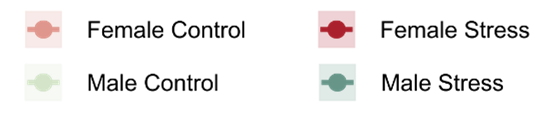

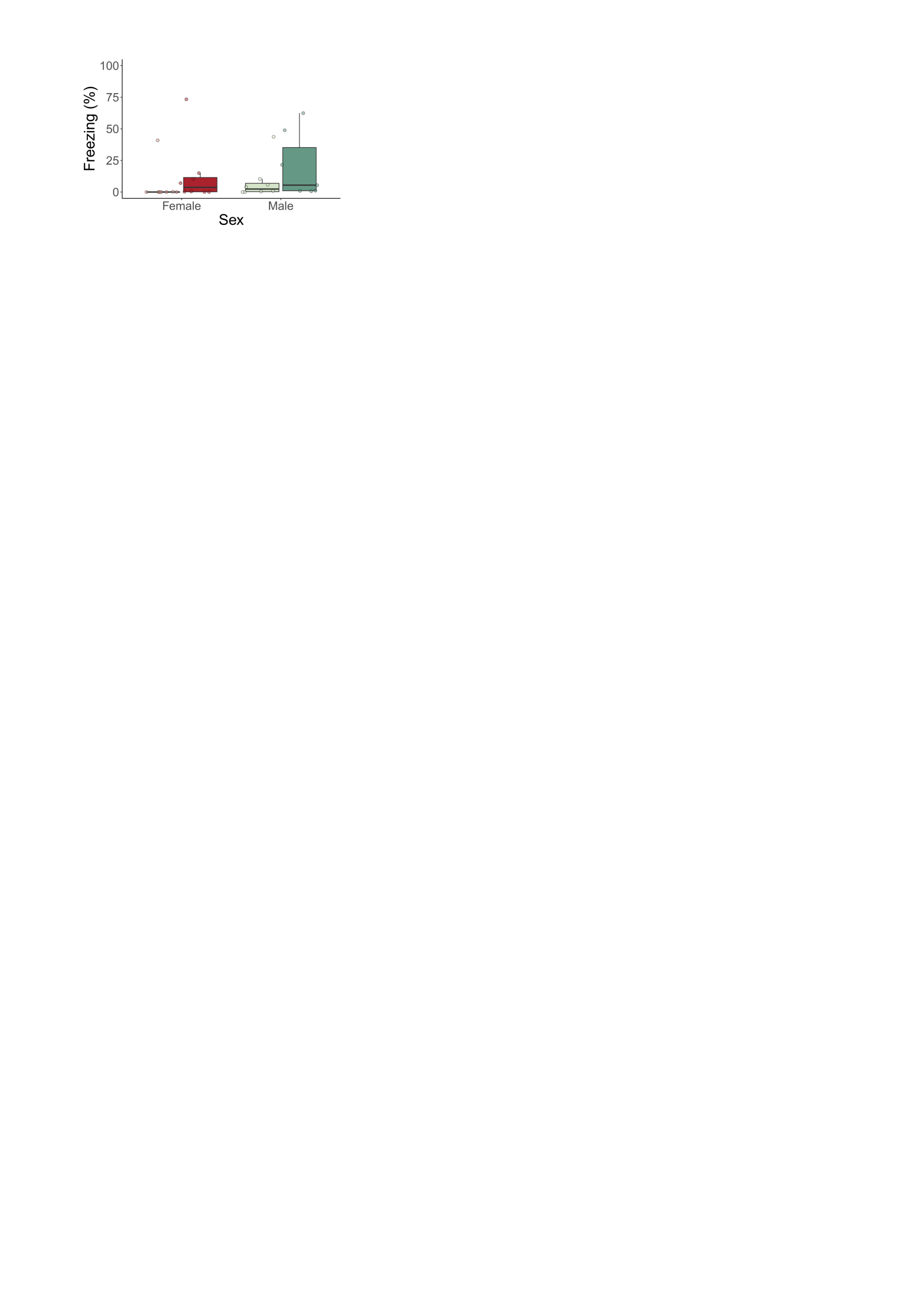

**Figure S1.** Experiment 1. Percentage of freezing during the 5-min acclimation period of day 2 avoidance acquisition in context A. There were no significant group nor sex differences.

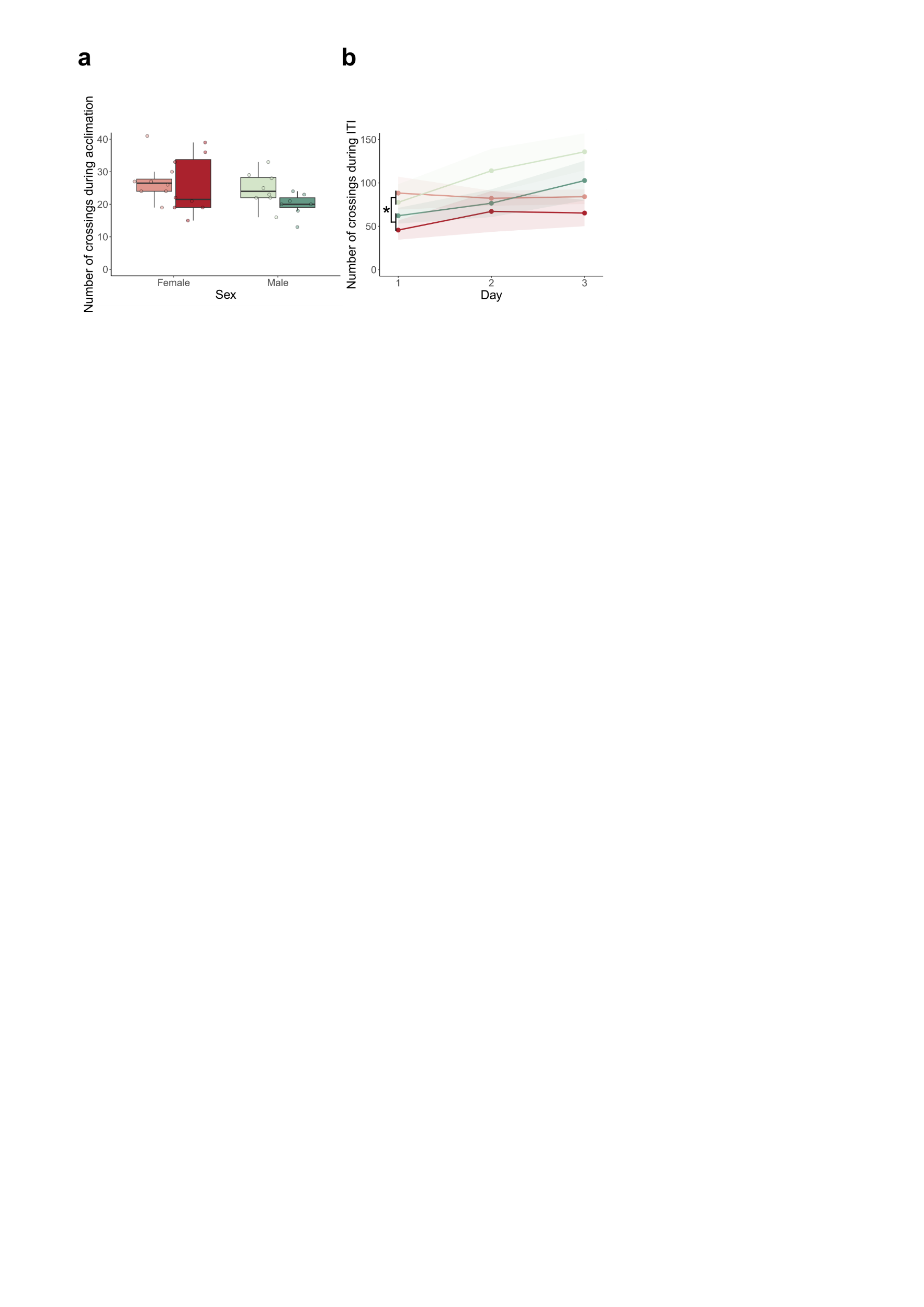

**Figure S2**. Experiment 1. Locomotion of rats during acquisition context A. **a.** Number of crossings during the acclimation period on the first day of avoidance acquisition in context A. Bold black lines represent the mean. All rats showed comparable numbers of crossings during this time period. **b.** Number of crossings across the intertrial intervals. Bold lines represent the mean, the surrounding shaded area represents the standard error of the mean. The mean number of crossings across the intertrial intervals (ITIs) was lower in stressed rats compared to control rats (F(1, 27) = 4.21, p = 0.05). We observed a main effect of Day (F(2, 54) = 3.8, p = 0.029), but post-hoc analyses did not identify significant pairwise comparisons.

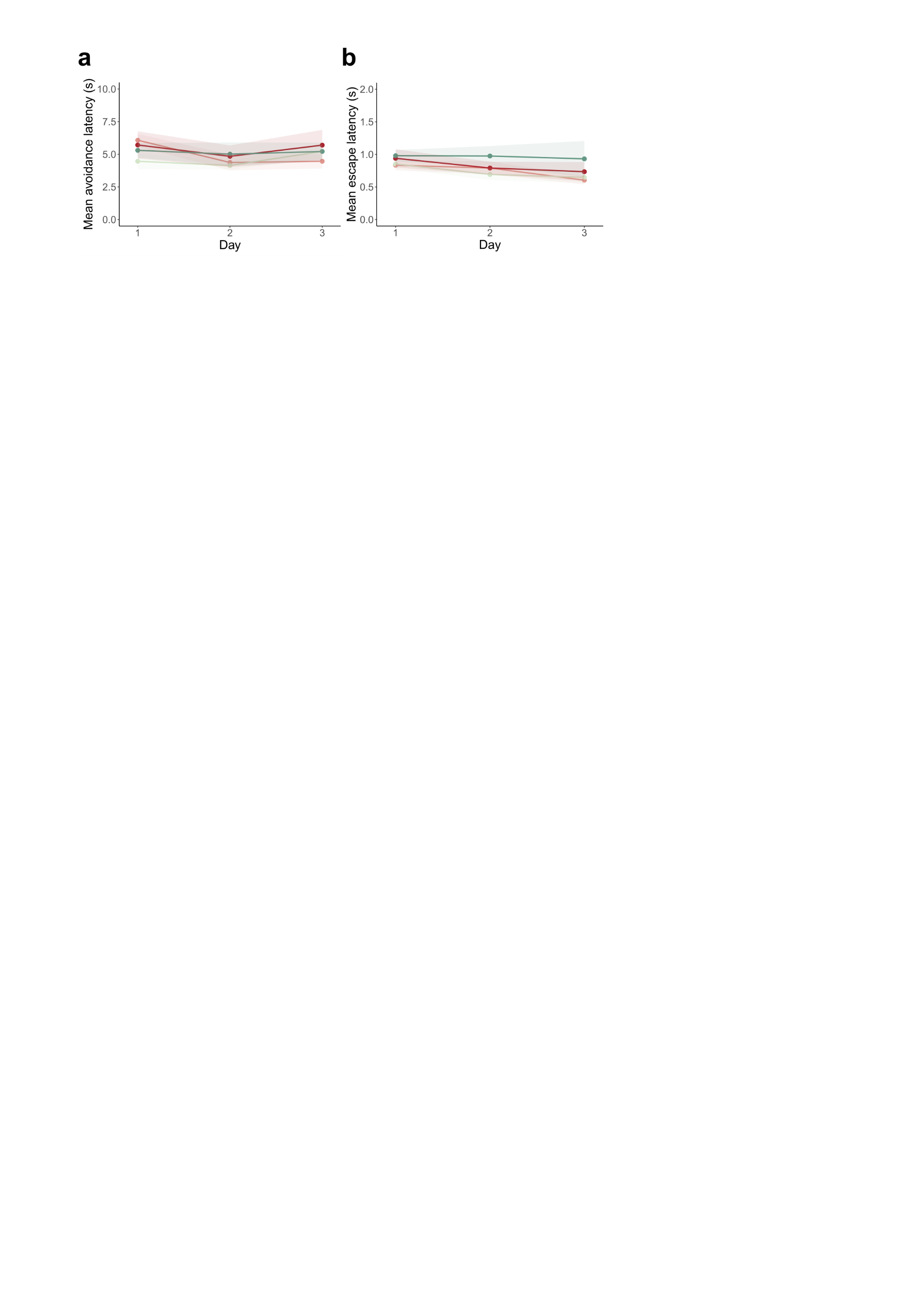

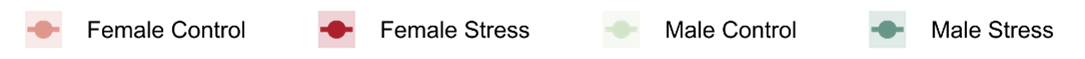

Figure S3. Experiment 1. Mean avoidance and mean escape latency in avoidance acquisition context A. Bold lines represent the mean, the surrounding shaded area represents the standard error of the mean. Note that avoidance latency has a maximum duration of 20 s and that escape latency has a maximum duration of 10 s. a. Mean avoidance latency did not differ between groups or sexes. b. Both groups and sexes showed similar escape latencies. We observed a main effect of Day (F(2, 52) = 3.93, p = 0.026), but post-hoc analyses did not identify significant pairwise differences.

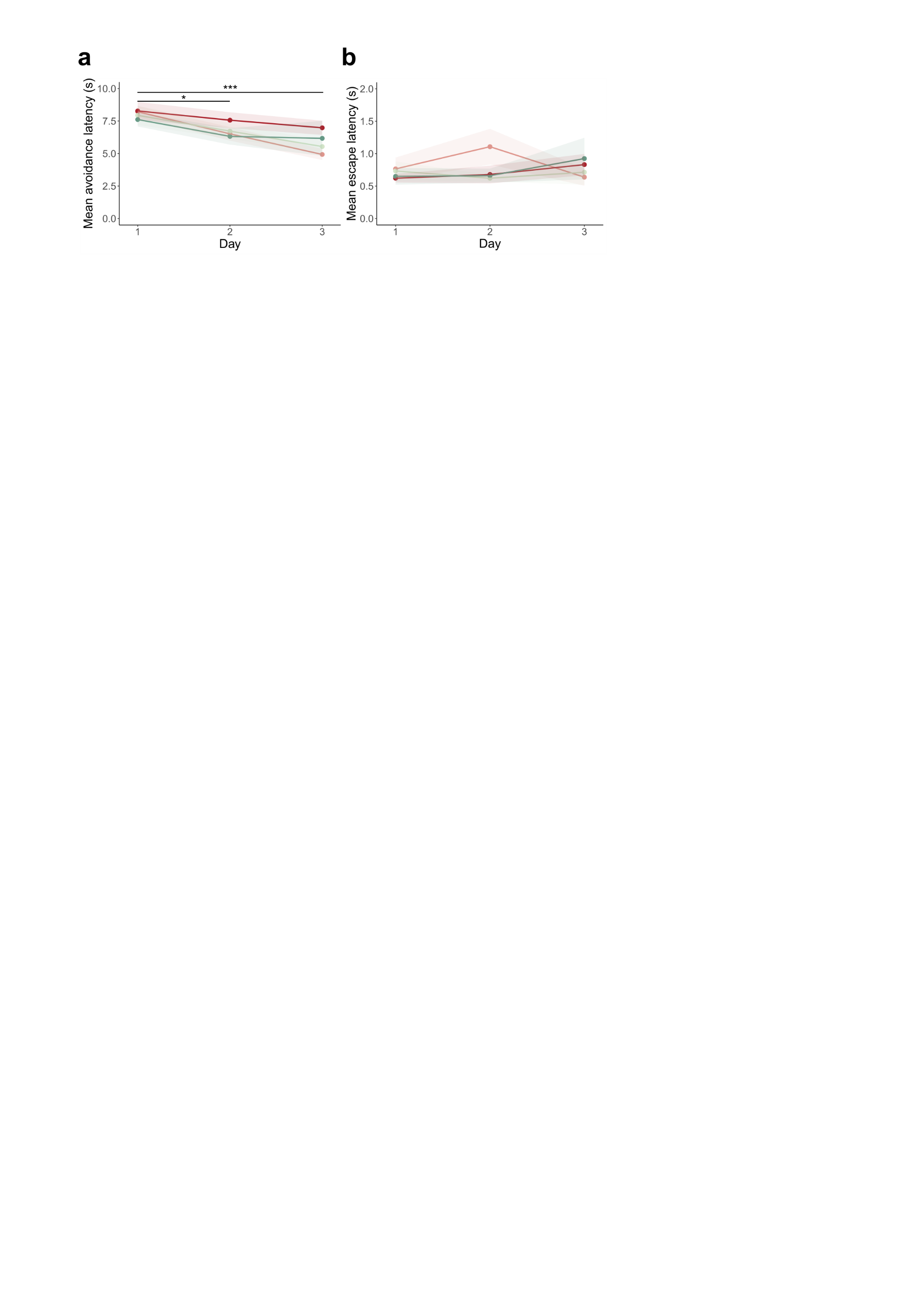

Figure S4. Experiment 1. Mean avoidance and mean escape latency in avoidance acquisition context B. Bold lines represent the mean, the surrounding shaded area represents the standard error of the mean. Note that avoidance latency has a maximum duration of 20 s and that escape latency has a maximum duration of 10 s. a. Rats showed similar avoidance latencies across groups and sexes. We observed a significant main effect of Day (F(2, 54) = 17.98, p < 0.001), where mean avoidance latency decreased from the first to the second day (t(81) = 2.68, p = 0.024) and from the first to the last day (t(81) = 4.58, p < 0.001) of acquisition in context B. b. Mean escape latency did not differ between groups or sexes.

Experiment 2

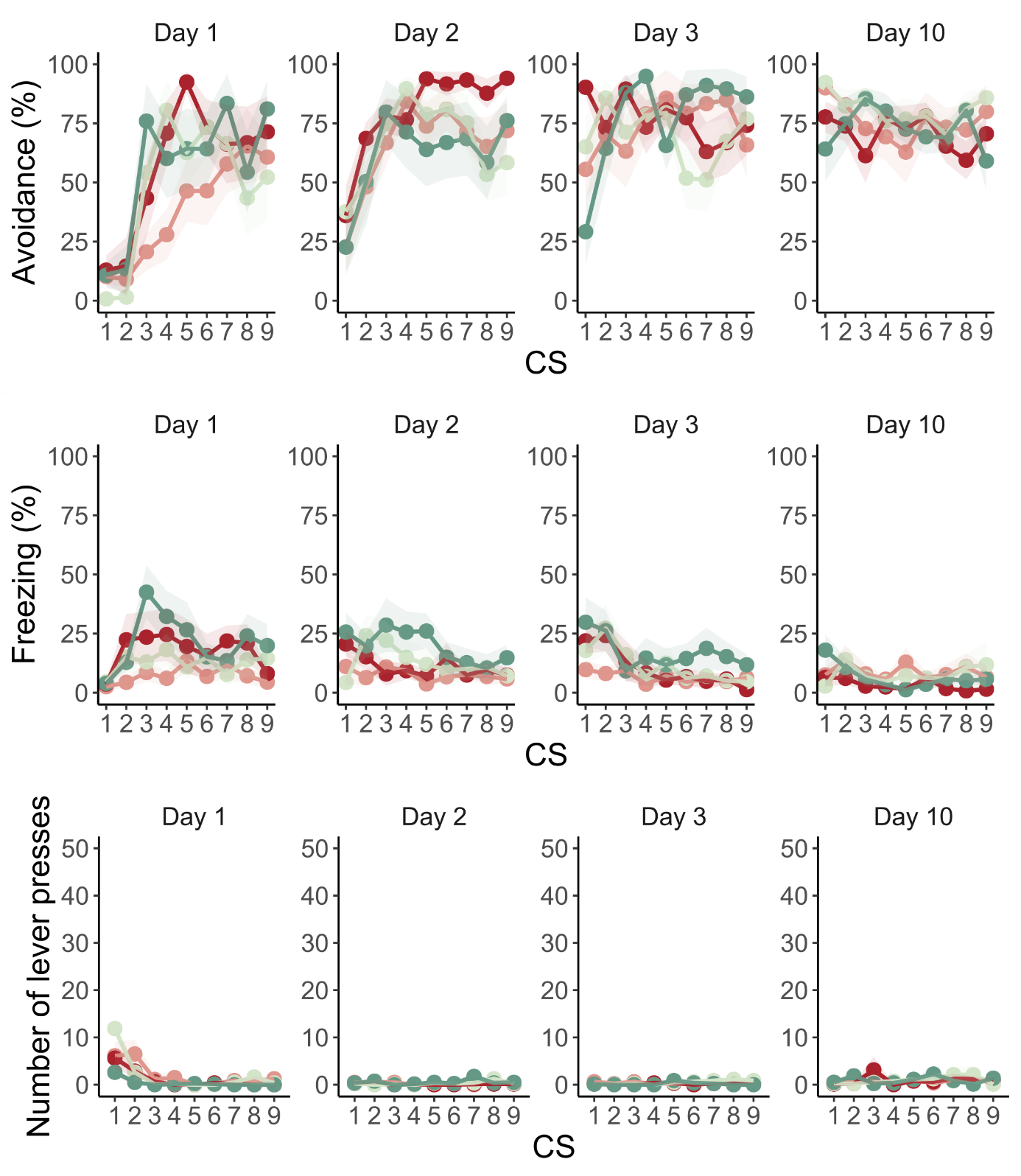

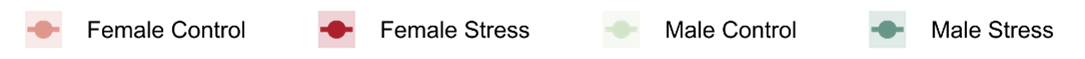

**Figure S5.** Experiment 2. Trial-by-trial data of acquisition days 1, 2, 3 and 10. The bold lines in the trial-by-trial plots represent the mean and the surrounding shaded area the standard error of the mean.

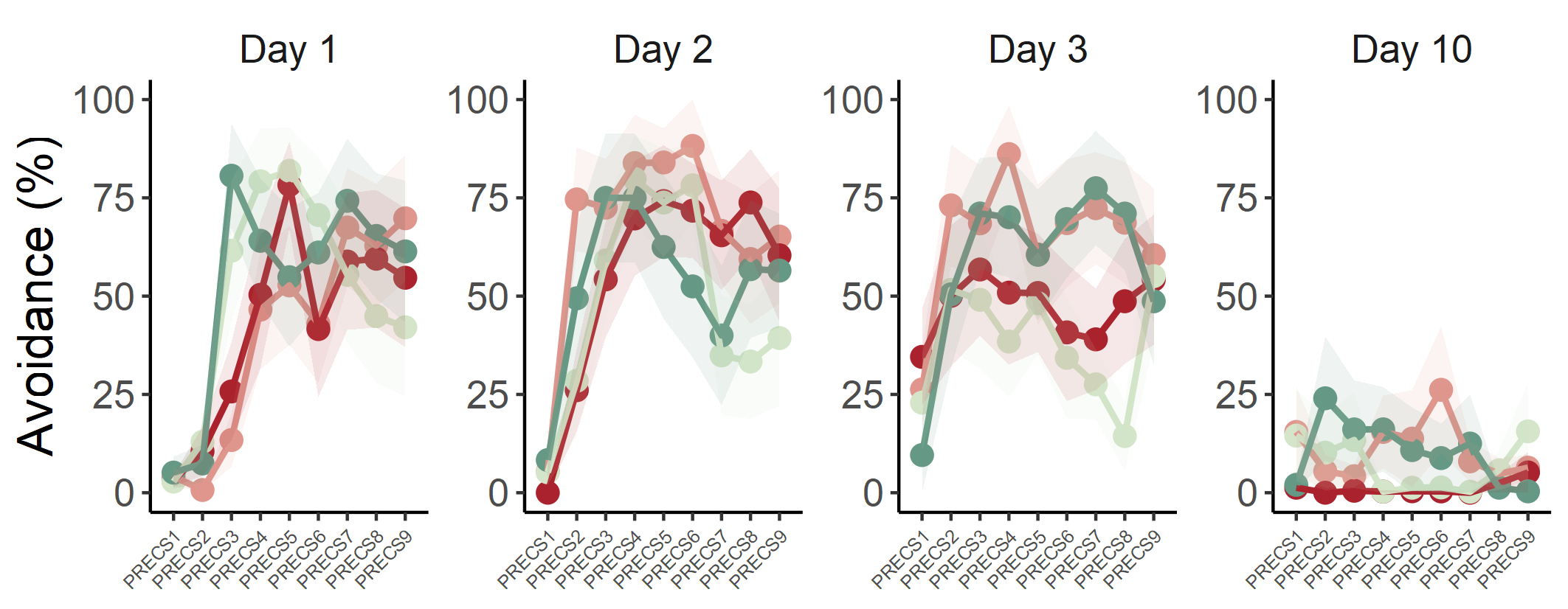

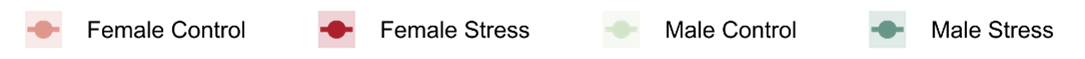

**Figure S6.** Experiment 2. Percentage of avoidance (i.e., time spent on the platform) during PRECS intervals (i.e., 1 minute before CS presentation) on day 1, 2, 3 and 10 of avoidance acquisition. The bold lines represent the mean of the percentage of avoidance per PRECS instance and the surrounding shaded area the standard error of the mean. We found a triple interaction between group, sex and day (F(3, 980) = 5.98, p < 0.001, ω_p_^2^ < 0.01), hence, we performed further analyses per day. On day 1, we found a significant interaction between PRECS interval and sex (F(8, 224) = 2.47, p = 0.014, ω_p_^2^ = 0.03), meaning that avoidance acquisition rates were different depending on sex. On day 3, we observed a significant interaction between group and sex (F(1, 28) = 4.62, p = 0.04, ω_p_^2^ =0.05). In both sexes, a significant difference between groups was observed (males: W = 1986.5, p = 0.013, r = 0.21 ; females: W = 3095, p = 0.037, r = 0.17). While male stressed animals showed higher avoidance levels (M = 58.7, SEM = 5.29) than controls (M = 38, SEM = 4.66), in females it was the opposite. Control female rats exhibited higher levels of avoidance (M = 65, SEM = 5.05) compared to stressed female rats (M = 47.3, SEM = 5.11).

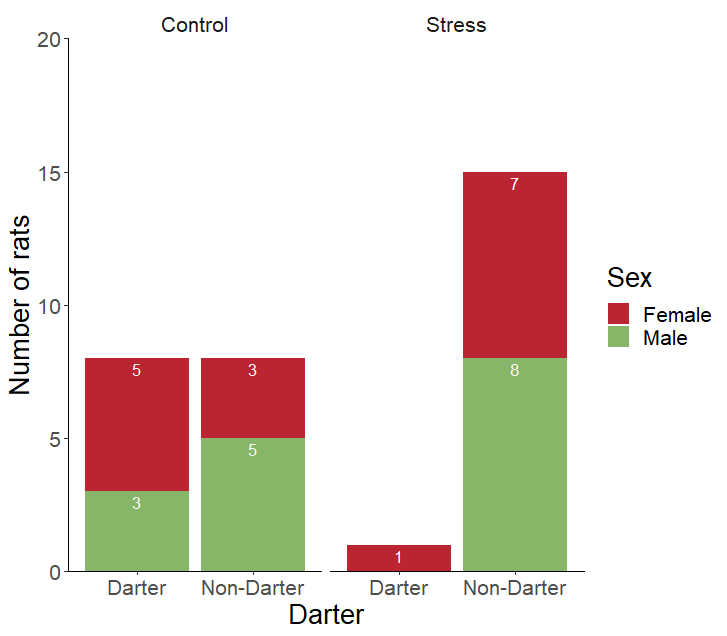

**Figure S7.** Experiment 2. Number of darter vs non-darter rats in control and stress conditions.

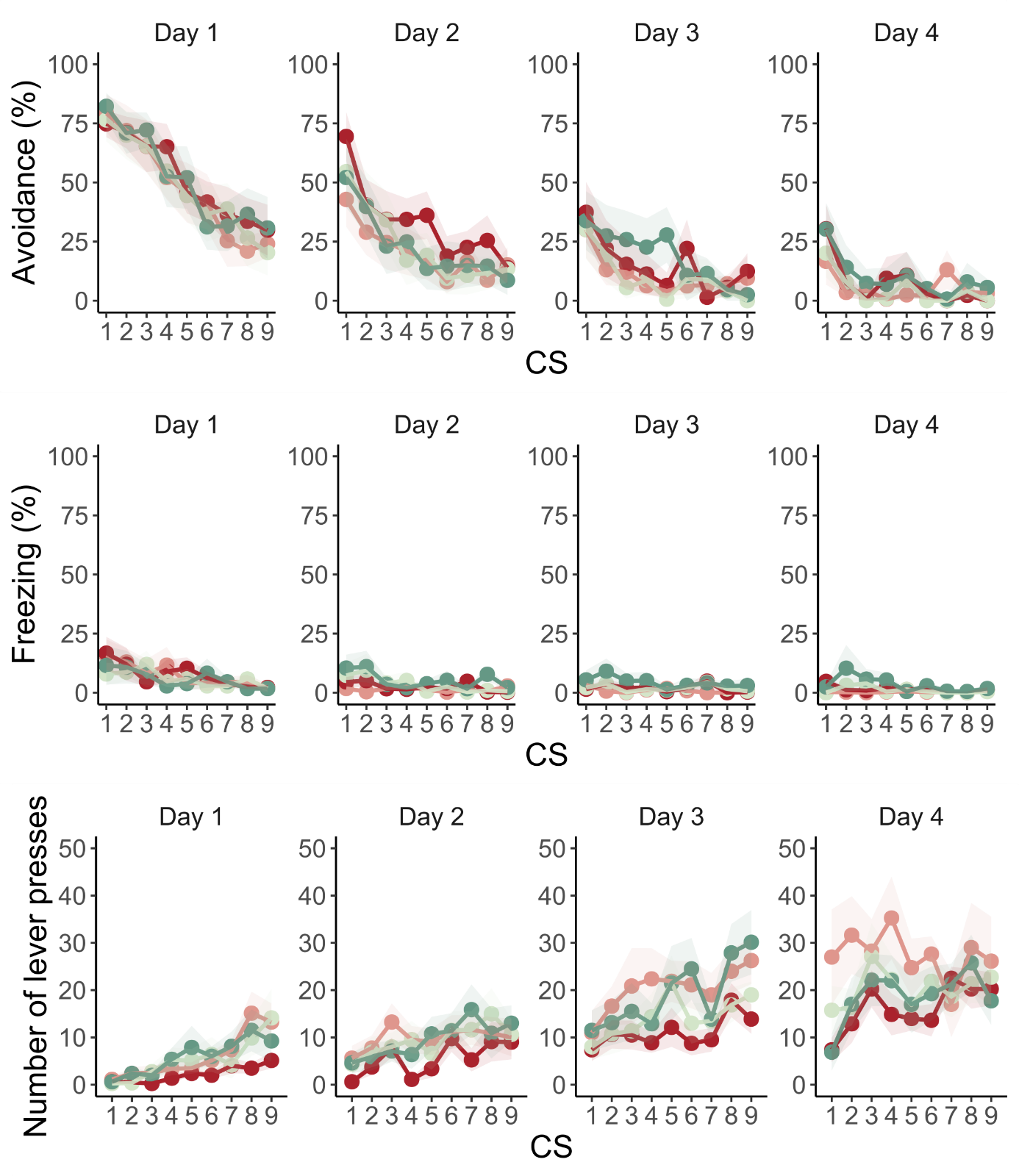

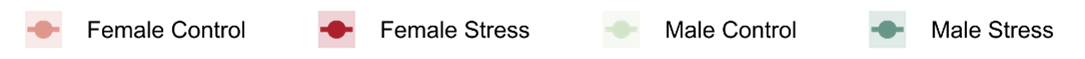

**Figure S8.** Experiment 2. Trial-by-trial data of all extinction sessions. The bold lines in the trial-by-trial plots represent the mean and the surrounding shaded area the standard error of the mean.

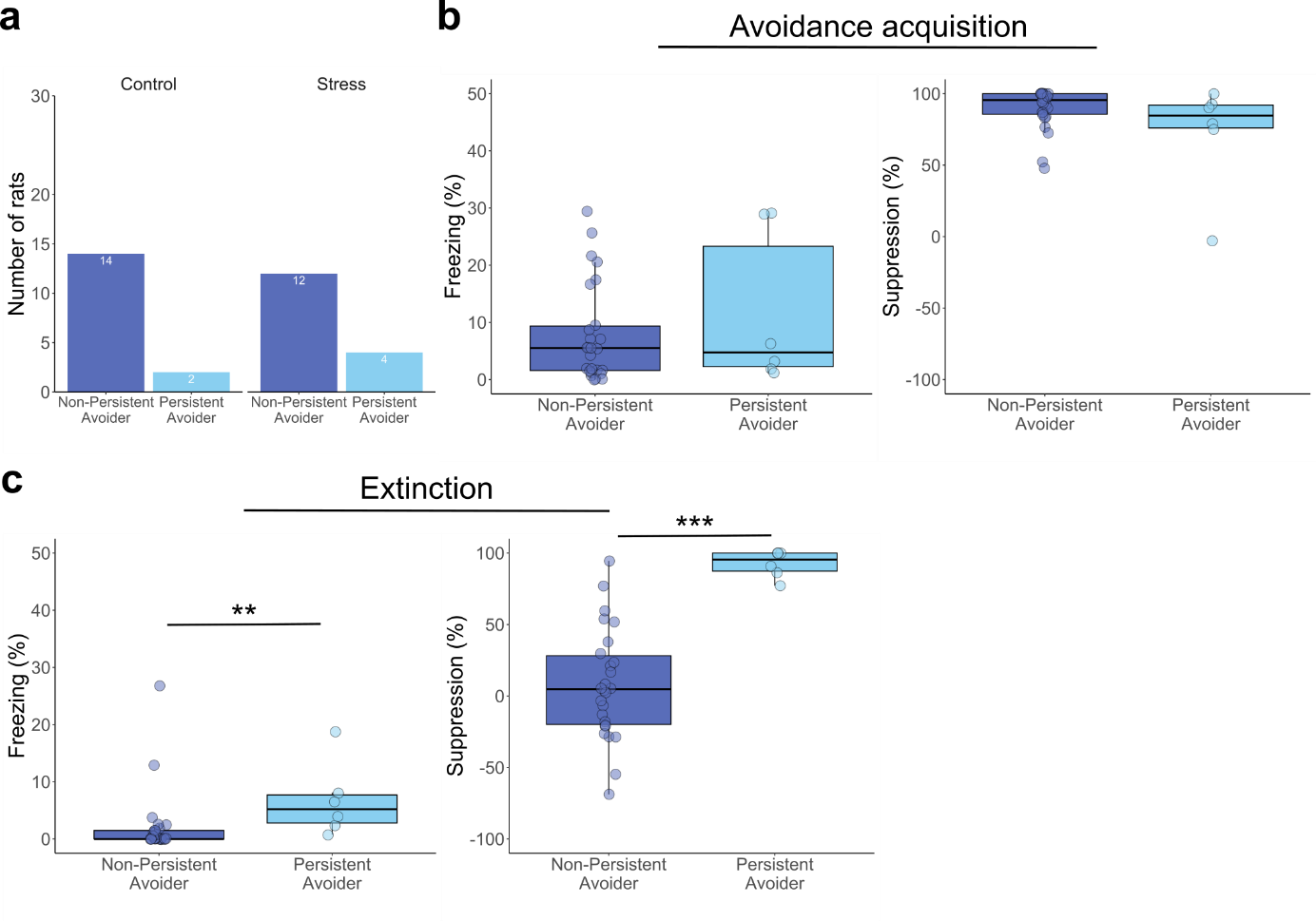

**Figure S9.** Experiment 2. Persistence of avoidance. **a**. Number of persistent avoider vs non-persistent avoider rats in control and stress conditions. **b.** Persistent and non-persistent avoider rats show similar levels of freezing and suppression of lever pressing during the last avoidance acquisition session. **c.** Persistent and non-persistent avoider rats show significant differences in freezing (W = 22, p = 0.005) and in suppression of lever pressing (W = 3, p < 0.001) on day 3 of extinction.

Supplementary Tables

Experiment 1

**Table S1.** Stress induction effects on body weight, including sex as factor - Experiment 1

| **Measured outcome** | **Statistical Test** | **Result** | **Effect size** |
| --- | --- | --- | --- |
| Weight change | Mixed ANOVA (Greenhouse-Geisser sphericity correction) | Sex: F(1, 27) = 24.41, **p < 0.001**  Group: F(1, 27) = 391.49, **p < 0.001**  Day: F(4.89,132.10) = 183.02, **p < 0.001**  S*G: F(1, 27) = 22.82, **p < 0.001**  S*D: F(4.89, 132.10) = 2.46, **p = 0.035**  G*D: F(4.89, 132.10) = 98.59, **p < 0.001**  S*G*D: F(4.89, 132.10) = 8.9, **p < 0.001** | ω_p_^2^ = 0.45 [0.21, 1]  ω_p_^2^ = 0.93 [0.89, 1]  ω_p_^2^ = 0.77 [0.74, 1]  ω_p_^2^ = 0.43 [0.19, 1]  ω_p_^2^ = 0.03 [0, 1]  ω_p_^2^ = 0.65 [0.59, 1]  ω_p_^2^ =  0.13 [0.05, 1] |
| Weight change | Effects by sex (males) - Mixed ANOVA (Greenhouse-Geisser sphericity correction) | Group: F(1, 13) = 273.72, **p < 0.001**  Day: F(3.09, 40.17) = 155.03, **p < 0.001**  G*D: F (3.09, 40.17) = 117.49, **p < 0.001** | ω_p_^2^ = 0.95 [0.89, 1]  ω_p_^2^ = 0.82 [0.77, 1]  ω_p_^2^ = 0.77 [0.71, 1] |
| Weight change | Simple effects by day (males): two-sample t-test with Bonferroni correction | Day 2 – Group: t(13) = 13.06, **p < 0.001**  Day 6 – Group: t(13) = 11.63, **p < 0.001**  Day 10 – Group: t(13) = 16.72, **p < 0.001** | d = 6.59 [4.28, 18.3]  d = 5.93 [4.34, 12.6]  d = 8.45 [6.67, 16.2] |
| Weight change | Effects by sex (females) - Mixed ANOVA | Group: F(1, 14) = 124.2, **p < 0.001**  Day: F(9, 126) = 62.55, **p < 0.001**  G*D: F(9, 126) = 21.31, **p < 0.001** | ω_p_^2^ = 0.89 [0.76, 1]  ω_p_^2^ = 0.67 [0.58, 1]  ω_p_^2^ = 0.33 [0.19, 1] |
| Weight change | Simple effects by day (females): two-sample t-test with Bonferroni correction | Day 2 – Group:  t(14) = 1.6365, **p = 0.011**  Day 6 - Group:  t(14) = 7.75, **p < 0.001**  Day 10 – Group: t(14) = 6.83, **p < 0.001** | d= 1.75 [0.78, 3.68]  d = 3.88 [2.94, 6.61]  d = 3.42 [2.43, 5.87] |

**Table S2.** Stress induction effects on acclimation freezing, acclimation crossings, ITI crossings, avoidance latencies and escape latencies - Experiment 1

| **Test phase** | **Measured outcome** | **Statistical Test** | **Result** | **Effect size** |
| --- | --- | --- | --- | --- |
| Acquisition context A day 2 | Percentage freezing during acclimation | Two-way non-parametric ANOVA | Sex: F(1, 27) = 1.73, p = 0.199  Group: F(1, 27) = 3.05, p = 0.092  S*G: F(1, 27) = 0.17, p = 0.684 | ω_p_^2^ = 0 [0, 1]  ω_p_^2^ = 0.03 [0, 1]  ω_p_^2^ = 0 [0, 1] |
| Acquisition context A day 1 | Number of crossings during acclimation | Two-way ANOVA | Sex: F(1, 27) = 3.05, p = 0.092  Group: F(1, 27) = 2.03, p = 0.166  S*G: F(1, 27) = 0.45, p = 0.506 | ω_p_^2^ = 0.06 [0, 1]  ω_p_^2^ = 0.03 [0, 1]  ω_p_^2^ = 0 [0, 1] |
| Acquisition context A | Number of crossings during ITIs | Mixed ANOVA | Sex: F(1, 27) = 2.96, p = 0.097  Group: F(1,27) = 4.21, **p = 0.05**  Day: F(2,54) = 3.8, **p = 0.029**  S*G: F(1, 27) = 0.01, p = 0.906  S*D: F(2, 54) = 2.02, p = 0.143  G*D: F(2, 54) = 0.01, p = 0.987  S*G*D: F(2, 54) = 0.81, p = 0.449 | ω_p_^2^ = 0.06 [0, 1]  ω_p_^2^ = 0.10 [0, 1]  ω_p_^2^ = 0.04 [0, 1]  ω_p_^2^ = 0 [0, 1]  ω_p_^2^ = 0.02 [0, 1]  ω_p_^2^ = 0 [0, 1]  ω_p_^2^ =  0 [0, 1] |
| Acquisition context A | Number of crossings during ITIs | Pairwise comparisons with Tukey correction | Day 1-2: t(81) = -1.31, p = 0.393  Day 1-3: t(81) = -2.26, p = 0.067  Day 2-3: t(81) = -0.95, p = 0.61 | d = -0.33 [-0.84, 0.18]  d = -0.58 [-1.09, -0.06]  d = -0.24 [-0.75, 0.27] |
| Acquisition context A | Mean avoidance latency | Mixed ANOVA | Sex: F(1, 27) = 0.31, p = 0.581  Group: F(1, 27) = 0.88, p = 0.356  Day: F(2, 54) = 1.84, p = 0.168  S*G: F(1, 27) = 0.01, p = 0.914  S*D: F(2, 54) = 1.06, p = 0.355  G*D: F(2, 54) = 0.15, p = 0.858  S*G*D: F(2, 54) = 1.07, p = 0.352 | ω_p_^2^ =  0 [0, 1]  ω_p_^2^ =  0 [0, 1]  ω_p_^2^ =  0.01 [0, 1]  ω_p_^2^ =  0 [0, 1]  ω_p_^2^ =  0 [0, 1]  ω_p_^2^ =  0 [0, 1]  ω_p_^2^ =  0 [0, 1] |
| Acquisition context A | Mean escape latency | Mixed ANOVA | Sex: F(1, 26) = 0.41, p = 0.529  Group: F(1, 26) = 2.39, p = 0.135  Day: F(2, 52) = 3.93, **p = 0.026**  S*G: F(1, 26) = 0.58, p = 0.453  S*D: F(2, 52) = 0.32, p = 0.726  G*D: F(2, 52) = 0.29, p = 0.75  S*G*D: F(2, 52) = 0.57, p = 0.571 | ω_p_^2^ =  0 [0, 1]  ω_p_^2^ =  0.05 [0, 1]  ω_p_^2^ =  0.03 [0, 1]  ω_p_^2^ =  0 [0, 1]  ω_p_^2^ =  0 [0, 1]  ω_p_^2^ =  0 [0, 1]  ω_p_^2^ =  0 [0, 1] |
| Acquisition context A | Mean escape latency | Pairwise comparisons with Tukey correction | Day 1-2: t(78) = 1.03, p = 0.56  Day 1-3: t(78) = 0.98, p = 0.124  Day 2-3: t(78) = 0.95, p = 0.61 | d = 0.26 [-0.25, 0.78]  d = 0.51 [-0.01, 1.03]  d = 0.25 [-0.27, 0.76] |
| Acquisition context B | Mean avoidance latency | Mixed ANOVA | Sex: F(1, 27) = 0.51, p = 0.480  Group: F(1, 27) = 1.02, p = 0.322  Day: F(2, 54) = 17.98, **p < 0.001**  S*G: F(1, 27) = 1.10, p = 0.294  S*D: F(2, 54) = 0.21, p = 0.811  G*D: F (2, 54) = 2.28, p = 0.112  S*G*D: F(2, 54) = 0.37, p = 0.693 | ω_p_^2^ =  0 [0, 1]  ω_p_^2^ =  0 [0, 1]  ω_p_^2^ =  0.19 [0.05, 1]  ω_p_^2^ =  0 [0, 1]  ω_p_^2^ =  0 [0, 1]  ω_p_^2^ =  0.02 [0, 1]  ω_p_^2^ =  0 [0, 1] |
| Acquisition context B | Mean avoidance latency | Pairwise comparisons with Tukey correction | Day 1-2: t(81) = 2.68, **p = 0.024**  Day 1-3: t(81) = 4.58, **p < 0.001**  Day 2-3: t(81) = 1.9, p = 0.144 | d = 0.68 [0.16, 1.2]  d = 1.17 [0.63, 1.7]  d = 0.48 [-0.03, 1] |
| Acquisition context B | Mean escape latency | Non-parametric mixed ANOVA | Sex: F(1, 26) = 0.7, p = 0.410  Group: F(1, 26) = 0.08, p = 0.774  Day: F(2, 52) = 2.11, p = 0.132  S*G, F(1, 26) = 1.02, p = 0.321  S*D: F(2, 52) = 0.58, p = 0.562  G*D: F(2, 52) = 1.91, p = 0.158  S*G*D: F(2, 52) = 1.61, p = 0.21 | ω_p_^2^ =  0 [0, 1]  ω_p_^2^ =  0 [0, 1]  ω_p_^2^ =  0 [0, 1]  ω_p_^2^ =  0 [0, 1]  ω_p_^2^ =  0 [0, 1]  ω_p_^2^ =  0 [0, 1]  ω_p_^2^ =  0 [0, 1] |

Experiment 2

**Table S3** Stress induction effects on body weight, including sex as factor - Experiment 2

| **Measured outcome** | **Statistical Test** | **Result** | **Effect size** |
| --- | --- | --- | --- |
| Weight change | Mixed ANOVA (Greenhouse-Geisser sphericity correction) | Sex: F(1, 28) = 8.673, **p = 0.006**  Group: F(1, 28) = 169.56, **p < 0.001**  Day: F(3.96, 110.96) = 91.31, **p < 0.001**  S*G: F(1, 28) = 4.47, **p = 0.044**  S*D: F(3.96, 110.96) = 12.42, **p < 0.001**  G*D: F(3.96, 110.96) = 94.12, **p < 0.001**  S*G*D: F(3.96, 110.96) = 4.41, **p = 0.02** | ω_p_^2^ = 0.2 [0.03, 1]  ω_p_^2^ = 0.85 [0.75, 1]  ω_p_^2^ = 0.46 [0.38, 1]  ω_p_^2^ = 0.1 [0.00, 1]  ω_p_^2^ = 0.1 [0.02, 1]  ω_p_^2^ = 0.47 [0.39, 1]  ω_p_^2^ = 0.03 [0.00, 1] |
| Weight change | Effects by sex (males) - Mixed ANOVA (Greenhouse-Geisser sphericity correction) | Group: F(1, 14) = 91.26, **p < 0.001**  Day: F(2.9, 40.61) = 87.87, **p < 0.001**  G*D: F(2.9, 40.61) = 70.17, **p < 0.001** | ω_p_^2^ = 0.85 [0.69, 1]  ω_p_^2^ = 0.56 [0.46, 1]  ω_p_^2^ = 0.51 [0.39, 1] |
| Weight change | Simple effects by day: two-sample t-test | Day 2 – Group: t(14) = 4.02, **p = 0.001**  Day 6 – Group: t(14) = 9.53, **p < 0.001**  Day 10 – Group: t(14) = 10.05, **p < 0.001** | d = 2.01 [0.76, 3.21]  d = 4.8 [2.74, 6.8]  d = 5.02 [2.92, 7.02] |
| Weight change | Effects by sex (females) - Mixed ANOVA | Group: F(1, 14) = 79.87, **p < 0.001**  Day: F(9, 126) = 18.7, **p < 0.001**  G*D: F(9, 126) = 30.01, **p < 0.001** | ω_p_^2^ = 0.83 [0.66, 1]  ω_p_^2^ = 0.29 [0.15, 1]  ω_p_^2^ = 0.40 [0.26, 1] |
| Weight change | Simple effects by day: two-sample t-test | Day 2 – Group: t(14) = 4.35, **p < 0.001**  Day 6 – Group: t(14) = 7.56, **p < 0.001**  Day 10 – Group: t(14) = 8.03, **p < 0.001** | d = 2.18 [0.89, 3.42]  d = 3.78 [2.06, 5.46]  d = 4.01 [2.22, 5.76] |

**Table S4.** Differences in defensive behaviors between darters and non-darters in the control group - Experiment 2.

| **Test phase** | **Measured outcome** | **Statistical test** | **Result** | **Effect size** |
| --- | --- | --- | --- | --- |
| Avoidance acquisition | Avoidance | Non-parametric mixed ANOVA | Day: F(2, 28) = 21.67, **p < 0.001**  Darter: F(1, 14) = 0.01, p = 0.917  D*D: F( 2, 28) = 0.85, p = 0.436 | ω_p_^2^ = 0.44 [0.24, 1]  ω_p_^2^ = 0 [0, 1]  ω_p_^2^ < 0.01 [0, 1] |
| Avoidance acquisition | Freezing | Non-parametric mixed ANOVA | Day: F(2, 28) = 1.54, p = 0.233  Darter: F(1, 14) = 0.23, p = 0.637  D*D: F( 2, 28) = 0.4, p = 0.677 | ω_p_^2^ < 0.01 [0, 1]  ω_p_^2^ = 0 [0, 1]  ω_p_^2^ = 0 [0, 1] |
| Avoidance acquisition | Suppression of lever pressing | Non-parametric mixed ANOVA | Day: F(2, 28) = 9.15, **p < 0.001**  Darter: F(1, 14) = 0.11, p = 0.743  D*D: F( 2, 28) = 0.31, p = 0.738 | ω_p_^2^ = 0.2 [0.03, 1]  ω_p_^2^ = 0 [0, 1]  ω_p_^2^ = 0 [0, 1] |
| Extinction | Avoidance | Non-parametric mixed ANOVA | Day: F(1, 14) = 48.71, **p < 0.001**  Darter: F(1, 14) = 1.11, p = 0.31  D*D: F(1, 14) = 0.18, p = 0.678 | ω_p_^2^ = 0.27 [0.06, 1]  ω_p_^2^ = 0.04 [0, 1]  ω_p_^2^ = 0 [0, 1] |
| Extinction | Freezing | Non-parametric mixed ANOVA | Day: F(1, 14) = 16.01, **p < 0.001**  Darter: F(1, 14) = 0.27, p = 0.605  D*D: F(1, 14) = 0.01, p = 0.905 | ω_p_^2^ = 0.04 [0, 1]  ω_p_^2^ < 0.01 [0, 1]  ω_p_^2^ = 0 [0, 1] |
| Extinction | Suppression of lever pressing | Mixed ANOVA | Day: F(1, 14) = 13.6, **p = 0.002**  Darter: F(1, 14) = 0.02, p = 0.897  D*D: F(1, 14) = 0.21, p = 0.657 | ω_p_^2^ = 0.23 [0.04, 1]  ω_p_^2^ = 0 [0, 1]  ω_p_^2^ = 0 [0, 1] |
